# A single-cell transcriptome atlas of the aging human and macaque retina

**DOI:** 10.1101/2020.07.17.207977

**Authors:** Wenyang Yi, Yufeng Lu, Suijuan Zhong, Mei Zhang, Le Sun, Hao Dong, Mengdi Wang, Min Wei, Haohuan Xie, Hongqiang Qu, Rongmei Peng, Jing Hong, Ziqin Yao, Yunyun Tong, Wei Wang, Qiang Ma, Zeyuan Liu, Yuqian Ma, Shouzhen Li, Chonghai Yin, Jianwei Liu, Chao Ma, Xiaoqun Wang, Qian Wu, Tian Xue

**Author notes:** These authors contributed equally (Wenyang Yi, Yufeng Lu, Suijuan Zhong, Mei Zhang). Corresponding authors: Tian Xue,; Qian Wu,; Xiaoqun Wang,; Mei Zhang.

## Abstract

The human retina is a complex neural tissue that detects light and sends visual information to the brain. However, the molecular and cellular processes that underlie aging primate retina remain unclear. Here, we provide a comprehensive transcriptomic atlas based on 119,520 single cells of the foveal and peripheral retina of humans and macaques covering different ages. The molecular features of retinal cells differed between the two species, suggesting the distinct regional and species specializations of the human and macaque retinae. In addition, human retinal aging occurred in a region- and cell-type- specific manner. Aging of human retina exhibited a foveal to peripheral gradient. *MYO9A*^−^ rods and a horizontal cell subtype were greatly reduced in aging retina, indicating their vulnerability to aging. Moreover, we generated a dataset showing the cell-type- and region- specific gene expression associated with 55 types of human retinal disease, which provides a foundation to understand the molecular and cellular mechanisms underlying human retinal diseases. Together, these datasets are valuable for understanding the molecular characteristics of primate retina, as well as the molecular regulation of aging progression and related diseases.

## Introduction

The human retina is a specialized light-sensitive tissue of neurons, glia, and nourishing blood vessels [1–3]. The retina has long served as a model system for developmental and functional studies of the central nervous system [4–7]. Different cell types in the retina, including rod and cone photoreceptors as well as bipolar (BCs), amacrine (ACs), horizontal (HCs), ganglion (GCs) and glial cells, are packed together into a tightly organized network that converts incoming light into electrochemical signals, which are then relayed to the brain for visual formation [1]. In addition to this complexity, primates, including humans and monkeys, possess a specialized fovea, which is absent in rodent models. The fovea, which is responsible for high-acuity vision, is enriched in cone photoreceptors that directly receive light and are supported by specific morphological Müller glia (MGs) [8]. The molecular differences between the foveal and peripheral retina are important for understanding human visual function. To date, multiple analyses in primates have demonstrated the region-specific retinal transcriptomes in developing and adult foveal and peripheral retina, suggesting distinct transcriptional regulations in the two regions [3, 9–14].

As people age, a deterioration in photoreceptor structure occurs in the human retina, particularly in the foveal region [15, 16]. Patients aged 60 or older display obvious reductions in the number of foveal photoreceptors, and a decline in color discrimination and vision sensitivity [17]. Moreover, they are at high risk for retinal diseases, such as age-related macular degeneration (AMD). To date, however, the key information on molecular changes in the human retina during the aging process at single-cell resolution is lacking. In particular, it is unclear whether different cell types in the human foveal and peripheral retina exhibit distinct molecular changes during aging. The incomplete understanding of the aging process and underlying complexities has restricted the development of therapeutic strategies to slow or reverse retinal aging, which could otherwise help to postpone or rescue age-related retinal diseases.

Nonhuman primate (NHP) models have been widely used to study human retinal development and visual function [18, 19]. Although human and macaque retina share many similarities, such as genetics, anatomy, physiology, and immunology, differences exist between them. For example, at birth, the monkey fovea appears to have more mature cone morphology than that of human, whereas the human fovea develops rapidly during infancy [20]. In addition, the cone subtype ratios are different cross the two species [21]. So far, detailed molecular and cellular signatures between the different cell types in human and macaque retina remain elusive. In particular, whether various cell types in the human and macaque retina exhibit different molecular changes following aging is largely unknown.

As the primate retina is a heterogeneous tissue containing various cell types, the transcriptional heterogeneity of retinal cells cannot be detected using conventional bulk RNA-sequencing (RNA-seq). In contrast, single cell RNA-sequencing (scRNA-seq) has the power to detect region- and cell-type-specific changes, especially for rare cell types. Also, scRNA-seq can decipher ligand-receptor crosstalk between cell-cell interactions [22]. To decipher the detailed molecular processes that accompany retinal aging, we mapped > 119,000 cell transcriptomes of the human and macaque (*Macaca mulatta*) retina from young to old stages and explored related gene transcriptional regulation using scRNA-seq and the bulk Assay for Transposase-Accessible Chromatin with high-throughput sequencing (bulk ATAC-seq). Our data revealed that although the human and macaque retina showed high similarity in major cell types, the molecular features were distinguishable between species. Rods showed the greatest interspecies variation. Although rods from humans and macaques could be divided into two subtypes by MYO9A expression, the proportion of subtypes varied between the species. Interestingly, MGs and cones not only exhibited regional transcription differences, they also interacted with each other by ligand-receptor regulation. Furthermore, aging in the human foveal retina occurred earlier than that of the peripheral retina, and *MYO9A*^−^ rods and one HC subtype showed greater vulnerability to aging. Finally, we generated a dataset showing the cell-type- and region-specific gene expression associated with 55 types of human retinal disease. Overall, our study provides a valuable data source for understanding the regional and species specializations of the human and macaque retina as well as the molecular regulation of aging progression and related diseases.

## Results

### Single-cell transcriptomes of primate retina

We collected single-cell transcriptomic profiles of 119,520 cells, including 38,558 from six healthy human samples (one 8-day old infant and, five adults aged between 35 and 87 years old) and 80,962 from five macaque samples (one 2-year old juvenile and four adults aged between 4 and 23 years old) (Supplementary Table 1), which covered progressive retinal aging, using droplet-based scRNA-seq (10x Genomics platform). Retinae were dissociated from 1.5-2.0 mm and 5 mm diameter pieces around the center of the foveal pit to represent foveal and peripheral cells, respectively (Fig. 1a-c, Supplementary Fig. 1a). In total, cell atlases of 45,231 foveal and 74,289 peripheral cells were collected. The sequencing depth was analyzed by the gene number per cell in each sample (Supplementary Fig. 1b). We performed unbiased clustering of cell profiles from the human retina with both foveal and peripheral regions and found 56 clusters (Supplementary Fig. 1c). Cells from the 8-day old infant generally clustered together and differed from cells from other samples. As human photoreceptors do not reach maturation until four years old [23], the cells in the 8-day old infant were not as mature as those of the adults (Supplementary Fig. 1c). Interestingly, some clusters showed regional specificity, e.g., Clusters 13, 22, and 51 showed foveal identity and Clusters 21, 32, and 43 showed peripheral identity (Supplementary Fig. 1c). With known cell-type markers, we grouped 56 clusters into 11 major classes, including six neuronal cell types (rods, cones, HCs, ACs, BCs, and GCs), three glial cell types (MGs, astrocytes and microglia), endothelial cells, and pericytes (Fig. 1b, Supplementary Fig. 1d-e, Supplementary Table 2). Interestingly, we found that cells from Human-Clusters 46 and 51 (which were rod and MG subgroups, respectively) showed similar transcriptomic profiles, with high expression of *OTX2* and *RLBP1* (Supplementary Fig. 1e). *OTX2* is a homeodomain transcription factor expressed in early and mature photoreceptors [6, 10, 13], and *RLBP1* is a marker for MGs [24]. These two markers have not been found co-expressed in retinal cells previously. To validate the existence of the subtypes, we performed immunofluorescence staining and observed that ∼3% of RLBP1^+^ cells also expressed OTX2 in the human adult retina, in both foveal and peripheral regions (Fig. 1d-e), indicating the existence of these cells in the adult human retina. Therefore, Clusters 46 and 51 are previously unidentified retinal cell types. In the macaque dataset, 68 clusters were classified, and 10 major cell types were annotated (Fig. 1c, Supplementary Fig. 1f-g, Supplementary Table 3). Compared to humans, astrocytes were not detected in the macaque retina (Fig. 1c, Supplementary Fig. 1f-g), consistent with previous scRNA-seq results from *M. fascicularis* [3]. Thus, we identified all major retinal cell types and OTX2^+^RLBP1^+^ cells in the primate retina by scRNA-seq analysis.

**Figure 1.**
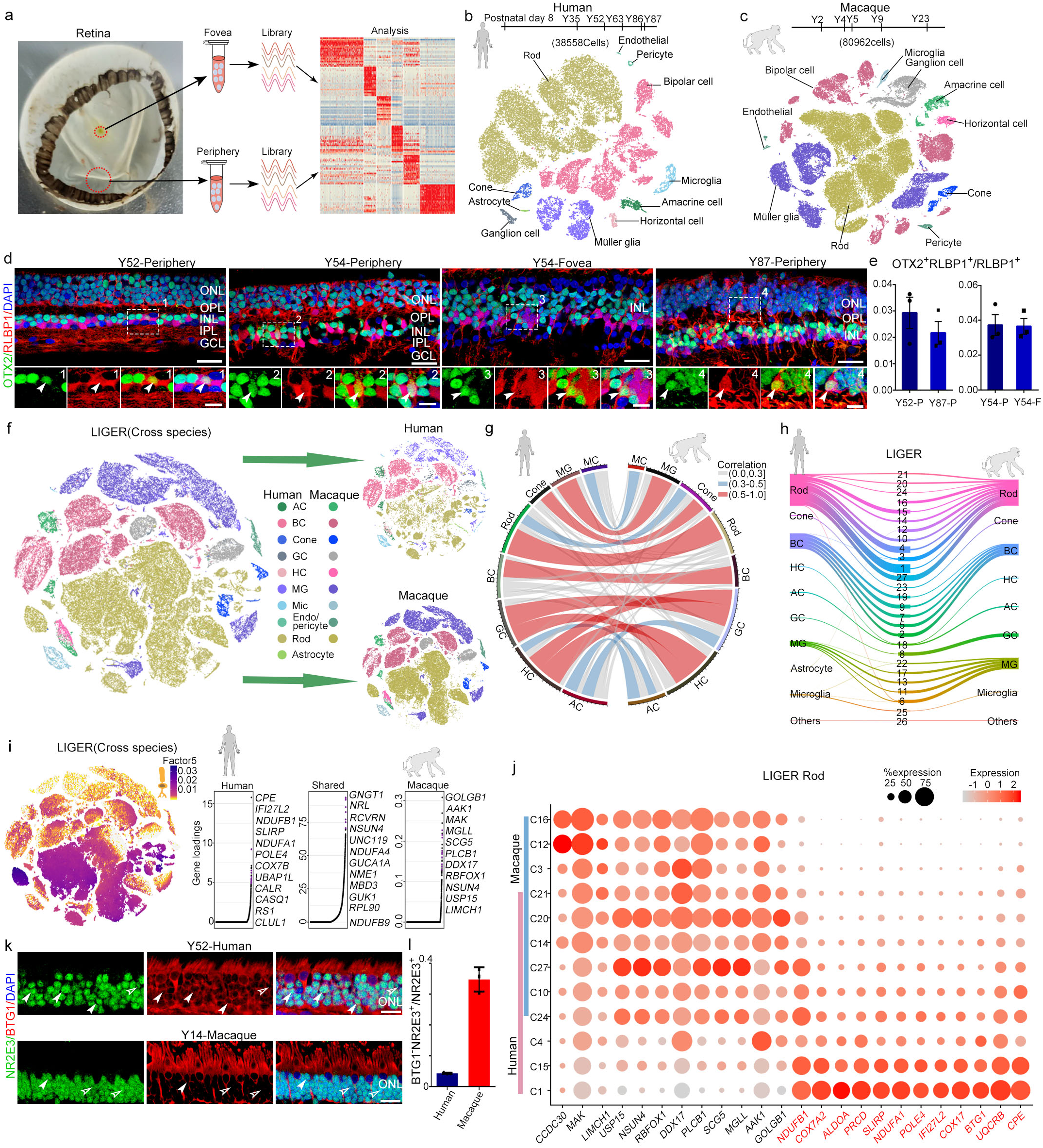
Cross-species transcriptome comparison of single-cell transcriptome profiles between human and macaque retina. (a) Experimental workflow for single-cell RNA-seq of human and macaque retina. Retinae were separated into foveal and peripheral samples for single-cell RNA-seq analysis. (b-c) t-SNE plot of single cells from human (b) and macaque (c) retina labelled with cell types. We obtained 38 558 human cells (samples: D8, Y35, Y52, Y63, Y86, and Y87) and 80 962 macaque cells (samples: Y2, Y4, Y5, Y9, and Y23). (d) Staining of OTX2 and RLBP1 of adult human retina in different age and region. Green: OTX2, Red: RLBP1, Blue: DAPI; bottom panel is higher-magnification view of top region boxed; Solid arrowheads indicate double positive cell. Scale bar, 25 μm (top), 10 μm (bottom). Experiments were repeated three times independently with similar results. (e) Bar charts showing proportion of OTX2^+^RLBP1^+^ cells out of RLBP1^+^ cells in human adult Y52 peripheral, Y87 peripheral, Y54 peripheral, and Y54 foveal retina, related to Fig. 1d. Data are means ± s.e.m. Each sample was counted from three different slices. (f) t-SNE visualization of 119 520 single cells analyzed by LIGER, color-coded by cell type. Top right is human and bottom right is macaque, with each dot representing a single cell (AC: amacrine cell, BC: bipolar cell, GC: ganglion cell, HC: horizontal cell, MG: Müller glia, Mic: microglia; Endo: endothelial). (g) Circos plot showing cross-species mapping between retinal cell types from humans and macaques. Correlation coefficients are expressed by width and color. Gray indicates correlation coefficient below 0.3, blue indicates correlation coefficient between 0.3 and 0.5, and red represents value greater than 0.5. (h) River plots comparing cell-type assignments for humans and macaques with LIGER joint clusters. (i) Cell factor loading values (left) and gene loading plots (right) of left loading dataset-specific and shared genes for factor 5. (j) Dot plot for specific genes related to factor 5 and differentially expressed genes (DEGs) of human and macaque rods in LIGER joint clusters, defined as rods. Macaque-specific genes are in black; human-specific genes are in red. Color of each dot shows average scale expression, and its size represents percentage of cells in cluster. (k) Immunostaining of NR2E3 and BTG1 in adult human and macaque retina. Solid arrowheads indicate NR2E3^+^BTG1^+^ cells, empty arrowheads indicate NR2E3+BTG1-cells. Scale bar: 15 μm. (l) Quantification of the proportion of BTG1^−^ NR2E3^+^ cells in adult human and macaque retina related to Fig. 1k. Each sample was counted from three different slices. Data are means ± s.e.m.

### Cross-species transcriptome comparison between human and macaque retina

To investigate the similarities and differences between the human and macaque retina, we integrated and analyzed the single-cell transcriptomic profiles to perform cross-species comparison using the LIGER algorithm [25], which can identify shared- or specific-“factors”, a cluster of genes that define specific cell types (Fig. 1f, Supplementary Fig. 1h). The major cell types showed high correlations between the two species, suggesting conservativeness between humans and macaques. In addition, GCs and HCs exhibited similar transcriptomes (Fig. 1g), which is possibly due to their close relationship in development [5, 7]. The LIGER joint clustering assignments indicated that rod clusters showed species-related differences, although they were well-mixed in the tSNE plots (Fig. 1f, h). Clusters 1, 4, and 15 were human-dominant, whereas Clusters 3, 12, and 16 were macaque-dominant (Fig. 1h). We next plotted the species-specific dimorphic genes derived from particular rod clusters (factor 5) in LIGER analysis (Fig. 1i). General rod marker genes (*GNGT1* and *NRL*) were shared, but species-specific genes for rod clusters in humans (*CPE*, *IFI27L2*, and *NDUFB1*) and macaques (*GOLGB1*, *AAK1*, and *MAK*) were also identified (Fig. 1i). Further analysis indicated that these genes also showed cluster specificity, consistent with species preference (Fig. 1j). For example, *CPE* (carboxypeptidase E), which is required for the maturation of photoreceptor synapse and its signal transmission to the inner retina [26], was highly expressed in human-specific clusters (Clusters 1 and 15) but minorly expressed in some shared clusters (Clusters 10 and 24) (Fig. 1j). To validate the species-specific marker (Fig. 1j), we stained BTG1, a cell cycle inhibitory gene [27], in both human and macaque retina (Fig. 1k and l). The results showed that BTG1 co-localized with NR2E3^+^ rods in outer nuclear layer (ONL) retina. However, the ratio of BTG1^−^ NR2E3^+^ cells to NR2E3^+^ cells was much lower in human retina, which is consistent with the scRNA-seq data. Although clusters of the cones and GCs from humans and macaques were closely matched by LIGER assignments (Clusters 23 and 8), we still found several species-specific genes in the cones (factor 18) and GCs (factor 6), respectively (Supplementary Fig. 1i). Collectively, our data demonstrated transcriptome conservativeness between the two species, but divergent gene expression was also discovered.

### Distinct subtypes of primate rod photoreceptors

Rods are the dominant cell type in the mammalian retina and are responsible for vision under low-light intensity [1, 28]. In total, we collected 16,686 rods in human retinal samples. As rods showed the highest interspecies differences by LIGER analysis, we next precisely analyzed rod subtypes. We first removed rod subclusters with more than 98% cells from the 8-day retinal sample to eliminate the influence of development. Based on analysis of differentially expressed genes (DEGs), human rods could be divided into two subgroups based on *MYO9A* expression (Fig. 2a-b). *CPE*, which was found to be dominantly expressed in humans by cross-species comparison (Fig. 1i), was highly expressed in *MYO9A*^−^ rods (Fig. 2c). In the human peripheral retina ONL, over 95% of cells were rods. Thus, we performed *MYO9A* RNAscope *in situ* hybridization (ISH) of the human peripheral retina and validated that a proportion of human rods expressed *MYO9A* (Fig. 2d), suggesting that human rods consist of *MYO9A^+^* and *MYO9A*^−^ cells. Consistent with human retina, macaque rods also consist of *MYO9A^+^* and *MYO9A*^−^ cells (Supplementary Fig. 2b). Moreover, subcluster of *MYO9A*^−^ in macaque retina was much lower than that in humans. To investigate the transcriptional regulation of *MYO9A*, bulk ATAC-seq was performed in human foveal and peripheral retinal samples (27, 54, and 86 years old). By searching transcription factor motifs from ATAC-seq peaks close to the *MYO9A* transcription start site (TSS), we found two potential OTX2 binding sites in *MYO9A* gene (Fig. 2e). Consistently, in humans, *OTX2* was generally expressed in the *MYO9A^+^* subcluster of rods (Fig. 2c, Supplementary Fig. 2a). Considering *OTX2* is a transcription factor controlling retinal photoreceptor cell fate [7], our results indicate that the expression of *MYO9A* might be regulated by *OTX2*. Accordingly, we observed both *OTX2*-positive and -negative cells in the human (Fig. 2f) and macaque retina (Supplementary Fig. 2c), similar to the division of *MYO9A*. From the scRNA-seq data, we found a similar *MYO9A^+^* to *MYO9A*^−^ rod ratio in the foveal and peripheral regions (Fig. 2g-h). As rods showed species specificity in their transcriptomes, we analyzed the macaque rods (36,904 cells) and observed that one subcluster (4.7% rods) was *MYO9A*^−^, which was much lower than that in humans (36.9%) (Fig. 2i-k) and consistent with ISH data (Supplementary Fig. 2b). Additionally, the distribution of *MYO9A*^−^ rods showed regional preference, with 98.6% of *MYO9A*^−^ rods located in the peripheral retina in macaques (Fig. 2l). Taken together, though the rods of humans and macaques could both be divided into two subtypes based on *MYO9A* expression, the proportion and location of *MYO9A*^−^ cells differed substantially between the two species.

**Figure 2.**
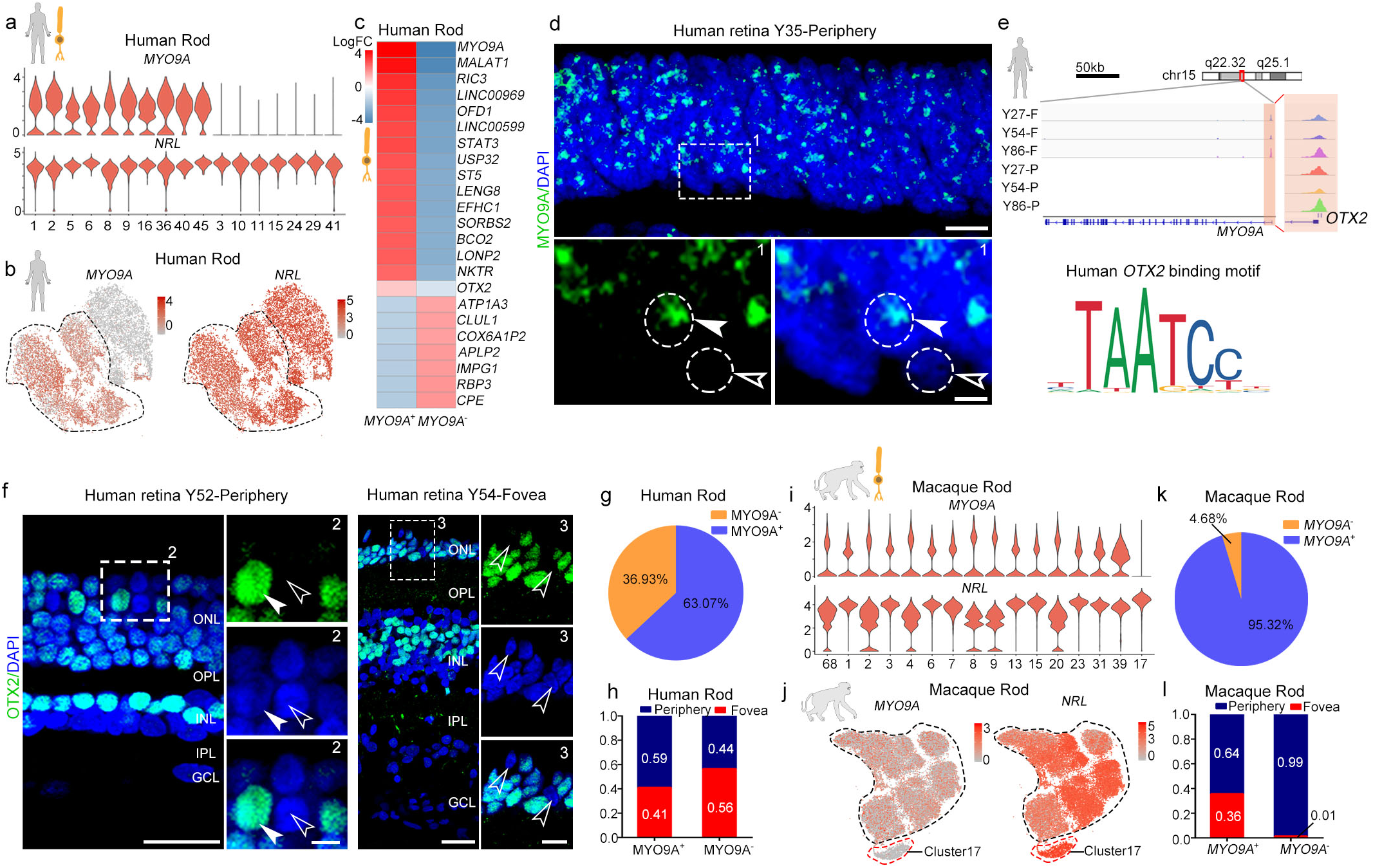
Distinct subtypes and differences in rods among primates. (a) Violin plots depicting expression of marker genes *MYO9A* and *NRL*, distinguishing 17 rod subclasses in human adult retina. (b) Visualization of expression of known marker genes *MYO9A* and *NRL* by t-SNE in human adult retina. Cells are colored according to gene expression levels (red, high; gray, low). (c) Heatmap illustrating log2 fold changes of differentially expressed genes (DEGs) between *MY09A*^+^ and *MYO9A*^−^ rod subclasses in human adult retina. (d) *In situ* RNA hybridization of *MYO9A* in adult human retina. Arrowheads and dotted circles show positive and negative cells, respectively; solid arrowhead indicates positive cell, empty arrowhead indicates negative cell. Blue, DAPI (nucleus marker). Scale bar, 10 μm (top), 5 μm (bottom). Experiments were repeated three times independently with similar results. (e) Normalized ATAC-seq profiles of *MYO9A* in fovea and periphery of Y27, Y54, and Y86 retina samples showing activation of *MYO9A*. Amplifying panel shows predicted *OTX2* binding sites (top). DNA binding motif of *OTX2* (bottom), identified in ATAC-seq peaks close to *MYO9A* transcription start site (TTS). (f) Immunostaining of OTX2 in adult human peripheral (left) and foveal retina (right). Solid arrowheads indicate positive cells; empty arrowheads indicate negative cells. Scale bar, 25 μm (left), 5 μm (right). Experiments were repeated three times independently with similar results. (g) Pie chart showing distribution of *MYO9A*^+^ and *MYO9A*^−^ subclasses in human adult retina samples. (h) Bar chart showing regional distributions of *MYO9A*^+^ and *MYO9A*^−^ rod subclasses in human retina samples. (i) Violin plots depicting expression of marker genes *MYO9A* and *NRL*, distinguishing 16 rod subclasses in macaque adult retina. (j) Visualization of expression of *MYO9A* and *NRL* by t-SNE in macaque retinae. Cells are colored according to gene expression levels (red, high; gray, low). (k) Pie chart showing distribution of *MYO9A*^+^ and *MYO9A*^−^ subclasses in macaque adult retina samples. (l) Bar chart showing regional distributions of *MYO9A*^+^ and *MYO9A*^−^ rod subclasses in macaque retina samples.

BCs are neurons that connect the outer to the inner layer in the retina, and play direct or indirect roles in transmitting signals from photoreceptors to GCs [1]. We found 17 and 18 BCs subclusters in humans and macaques, respectively, and categorized these subclusters into 12 and 11 known types in humans and macaques, respectively (Supplementary Fig. 2d-f). The 11 types of *M. mulatta* BCs were consistent with previous findings in *M. fascicularis* [3]. Similar to *M. fascicularis*, the Blue Bipolar (BB) and Giant Bipolar (GB) types were grouped together due to their transcriptional similarity in *M. mulatta*, whereas BB and GB types of human BCs could be separated by the transcriptome with distinct molecular markers (Supplementary Fig. 2e). However, the BC subtypes were generally conserved between humans and macaques (Supplementary Fig. 2g).

### Regionally different transcriptomes of primate MGs and cones

The foveal retina is cone-dominated, whereas the peripheral retina is rod-dominated. MGs are particularly important in the fovea because the foveola center is formed only by cone and MG processes [29]. To investigate the region-specific features of MGs and cones, we compared their single-cell transcriptomes in the foveal and peripheral retina. MGs exhibited remarkable regional specificity (Fig. 3a). Generally, most of foveal MGs were cells in Clusters 13 and 22 (Fig. 3a) and exhibited high *TRH*, *FGF9, DIO2, CYP26A1, RGR*, and *HTRA1* expression (Fig. 3b, Supplementary Fig. 3a-b). The majority of MGs in Clusters 21, 32, and 43 were located in peripheral retina and exhibited high *NPVF*, *MT3*, *MT2A*, and *GPX3* expression (Fig. 3b, Supplementary Fig. 3a-b). Immunostaining of NPVF and RLBP1 in the human retina validated the scRNA-seq results, indicating that *NPVF* was a marker gene of peripheral MGs in humans (Fig. 3c). In addition, immunostaining and ISH also revealed the foveal MG specific expression of TRH (thyrotropin releasing hormone) and DIO2 (iodothyronine deiodinase 2) respectively (Fig. 3d-e). We performed weighted gene co-correlation network analysis (WGCNA) of MG regional DEGs and found several foveal and peripheral MG-specific modules (Supplementary Fig. 3c-d). For the top foveal module, we performed network analysis and found that *TRH, RGR, DIO2,* and *CYP26A1* were also highly enriched (Fig. 3f). *CYP26A1* is a retinoic acid-metabolizing enzyme, with foveal-specific expression validated here (Supplementary Fig. 3e). *CYP26A1* has been identified as a DEG of the foveal retina during human development [30], which functions in rod-free zone generation [31, 32]. Interestingly, the expression level of *RGR* (retinal G protein-coupled receptor), another foveal MG-specific gene, is higher in the foveal MGs. *RGR* encodes a crucial factor for the photic visual cycle of cone visual pigment regeneration, mutations of which are associated with retinitis pigmentosa (RP) [33–35]. The cone visual cycle of regenerating 11-*cis* retinal is dependent on both retinal pigment epithelium (RPE) and MGs [34, 36]. Thus, the higher *RGR* expression in the foveal MGs might meet the high demand of cone visual pigment regeneration in the cone-dominated fovea. Additionally, *HTRA1*, a risk factor gene of AMD, exhibited high expression in the fovea MGs, consistent with the fovea being more vulnerable in AMD [37, 38]. Together, our results indicated that MGs in the fovea, as a type of supporting glial cell, expressed specific genes for facilitating cone photoreceptor regeneration, specification, and functional maintenance. We next analyzed the MGs of macaques, which also showed regional differences in terms of cell types and gene expression patterns (Supplementary Fig. 3f-g). Comparing the DEGs between the foveal and peripheral MG across both species, several genes showed similar regional specificity, such as *TRH, DIO2, CYP26A1, RGR, GPX3,* and *RPL7*, whereas gene expression preference of *NPVF, MT1F,* and *JUNB* was not conserved (Fig. 3g).

**Figure 3.**
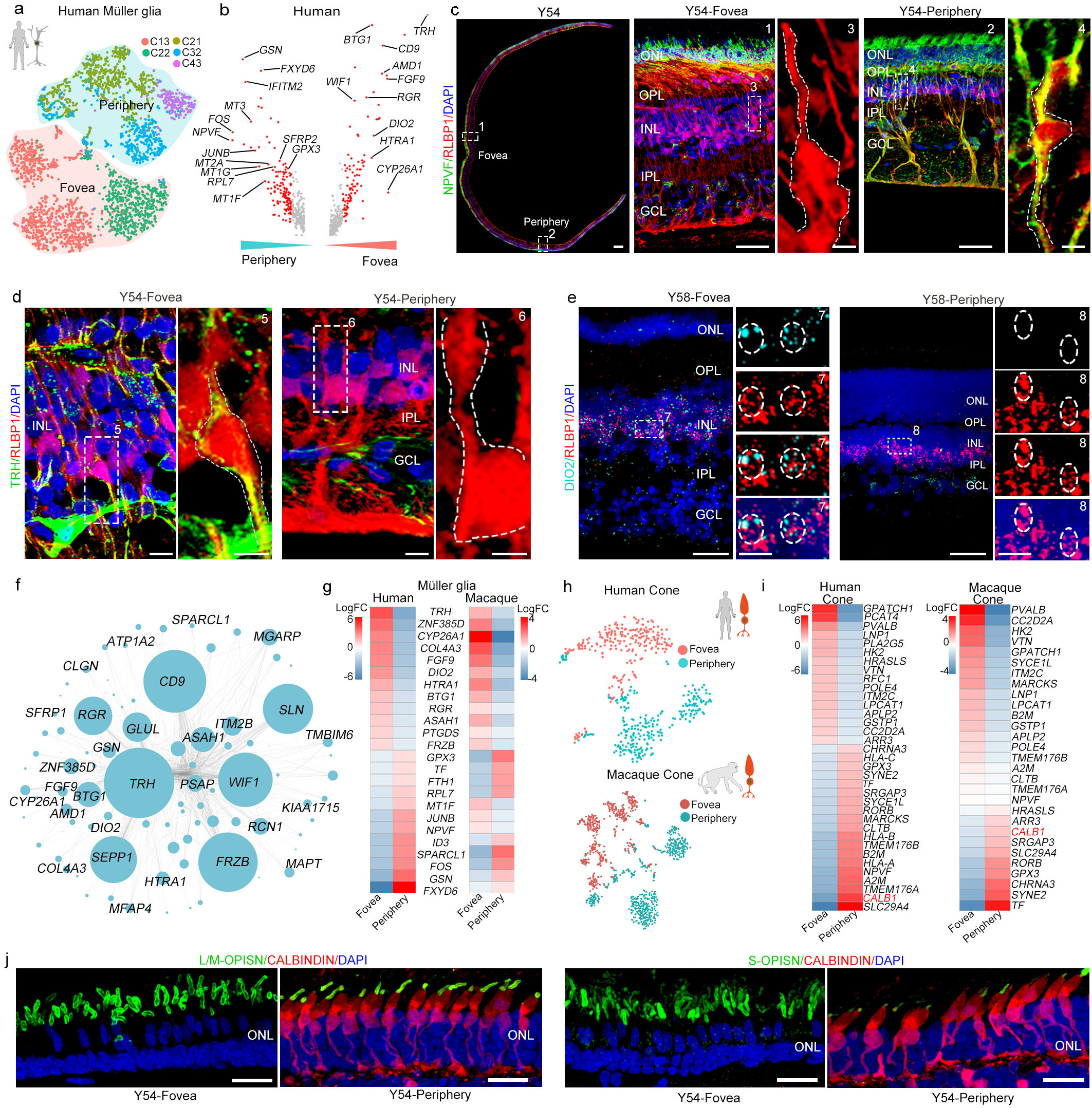
Regionally different transcriptomes of primate MG and cones. (a) t-SNE plot of MG distinguished by different subclasses (dots, individual cells; color, subclasses; color contours, regions). (b) Volcano plot of DEGs between human foveal and peripheral MG. Red dots show average log2 fold changes >0.5. (c) Immunostaining of RLBP1 and NPVF in Y54. Middle panel is higher magnification view of fovea in left-most panel; right panel is higher-magnification view of periphery in left-most panel. Scale bar of Y54, 300 μm, 50 μm (middle), 5 μm (middle on right), 50 μm (right), 5 μm (right-most). Experiments were repeated three times independently with similar results. (d) Immunostaining of RLBP1 and TRH in Y54 human retina, Scale bar, 10 μm (left), 5 μm (right). Experiments were repeated three times independently with similar results. (e) *In situ* RNA hybridization of *DIO2* and *RLBP1* in Y58 human retina. Blue, DAPI (nucleus marker). Scale bar, 50 μm (left), 10 μm (right). Experiments were repeated three times independently with similar results. (f) Network analysis of blue module genes. (g) Heatmap illustrating log2 fold changes of DEGs related to human foveal and peripheral Müller glia in human and macaque adult retina. (h) t-SNE plot of human (top) and macaque (bottom) cones based on regions (dots, single cell; color, regions). (i) Heatmap showing log2 fold changes in DEGs between foveal and peripheral cones in human (left) and macaque (right) adult retina. (j) Immunostaining of S-OPSIN/CALBINDIN and L/M-OPSIN/CABINDIN in Y54. Scale bar, 25 μm. Experiments were repeated three times independently with similar results.

Interestingly, cone photoreceptors also exhibited remarkable region-specificity in both humans and macaques (Fig. 3h). The thyroid hormone (TH) receptor, THRβ (human thyroid hormone receptor beta), is expressed in cones and also serves as a transcription factor [39]. Our ATAC-seq results showed that the potential binding sites of THRβ were found in open chromatin close to the TSS of several foveal-enriched genes (*RFC1, POLE4,* and *APLP2*) (Fig. 3i, Supplementary Fig. 3h). DIO2 converts the inactive TH thyroxine (T4) in blood into the active TH triiodothyronine (T3) [40]. Of note, we also found *DIO2* to be highly expressed in foveal MGs (Fig. 3b, e), indicating that foveal MGs may regulate the region-specific transcriptome of cones. The dosage of TH has been reported to play a role in specifying cone subtypes in human retinal organoids [41]. Here, our data demonstrated the MGs may modulate region-specific gene expression by TH signaling in foveal cones in adulthood. *CALB1* was one of the markers for peripheral cones, as determined by scRNA-seq data analysis (Fig. 3i). To confirm this finding, we carried out immunostaining of cone OPISN and CALBINDIN (encoded by gene *CALB1*) in human retina and found that CALBINDIN was only expressed in cones (both L/M and S cones) located in the peripheral retina (Fig. 3j, Supplementary Fig. 3i). Taken together, both MGs and cones displayed distinct regional gene expression profiles and MGs may modulate region-specific gene expression to facilitate cone photoreceptor generation, specification and functional maintenance by cell-cell communication.

### Molecular classification of retinal HCs in primates

HCs are located in the inner nuclear layer (INL) of the vertebrate retina, where they interconnect laterally with photoreceptors [42]. Here, the human HCs clustered into classical H1 (*LHX1^+^ PCP4^+^*) and H2 (*ISL1^+^ CALB1^+^*) subtypes (Fig. 4a-b, Supplementary Fig. 4a). Interestingly, H1 and H2 cell type showed regional preference, i.e., 92.0% of foveal HCs were H1 type and 60% of peripheral HCs were H2 type (Fig. 4c). Immunostaining of CALBINDIN/ONECUT2, and ISH of *LHX1* and *ISL1* in the human retina confirmed the scRNA-seq results, illustrating that most of human foveal HCs were H1 (Fig. 4d-f). The macaque HCs were also classified as H1 and H2 types (Fig. 4g, Supplementary Fig. 4b). Compared to the 92.0% in humans, 60.0% of macaque foveal HCs were H1 type (Fig. 4h). Although most DEGs between H1 and H2 showed similar expression across the two primate species, some human H1-enriched genes, e.g., *FILIPL1, CMSS1,* and *PCDH9*, were highly expressed in macaque H2 cells, whereas some human H2-enriched genes, e.g., *SYNPR, MGARP,* and *AKR1B1,* were highly expressed in macaque H1 cells (Fig. 4i, Supplementary Fig. 4c), indicating that distribution and gene expression were slightly different across the H1/H2 cell types of both species.

**Figure 4.**
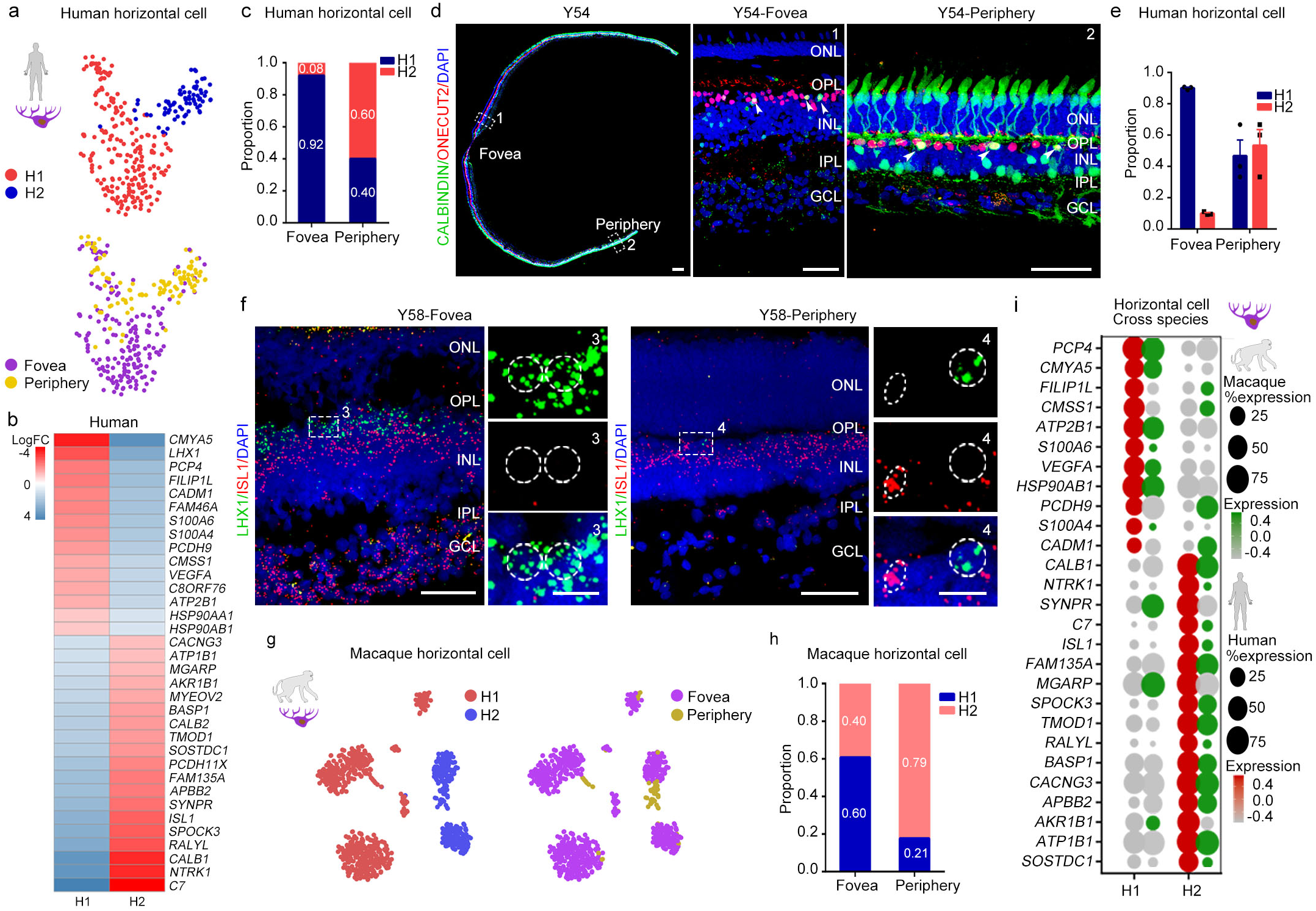
Molecular properties of retinal horizontal cells in primates. (a) t-SNE plot of human horizontal cells, colored by subclass (top) and region (bottom). Each dot is a single cell. (b) Heatmap showing log2 fold changes in DEGs between H1 and H2 horizontal cells in human adult retina. (c) Proportion of human horizontal cell subclasses in foveal and peripheral regions. (d) Immunostaining of CALBINDIN and ONECUT2 in Y54. Middle panel is higher-magnification view of fovea in left-most panel; right panel is higher-magnification view of periphery in left-most panel. Solid arrowheads indicate double positive cell. Scale bar of Y54, 300 μm, 50 μm (middle), 50 μm (right). Experiments were repeated three times independently with similar results. (e) Quantification of proportion of H1 and H2 subclasses in foveal and peripheral human adult retina related to Fig. 4d. Each sample was counted from three different slices. Data are means ± s.e.m. (f) *In situ* RNA hybridization of *LHX1* and *ISL1* in Y58 human retina. Blue, DAPI (nucleus marker). Scale bar, 50 μm (left), 10 μm (right). Experiments were repeated three times independently with similar results. (g) t-SNE visualization of macaque horizontal cells, color-coded by subclass (left) and region (right). Each dot represents a single cell. (h) Proportion of macaque horizontal cell subclasses in foveal and peripheral regions. (i) Dot plot showing H1- and H2-specific marker genes in human (red) and macaque (green) adult retina. Size of each dot represents percentage of cells in each cluster (red/green, high expression; gray, low expression).

### Aging patterns of human retina

Aging affects retinal function and structure physiologically and pathologically. To clarify the cellular and molecular changes during aging, we systematically studied human retinal samples (between 35 to 87 years old). We categorized the five samples in three groups (35Y, adult group; 52Y and 63Y, mid-age group; 86Y and 87Y, aging group). We first focused on whether the genes continuously increased or decreased from the adult to aging groups (Fig. 5a-e). In total, the expression of 87 genes was up-regulated. Gene ontology (GO) analysis indicated these genes may play roles in response to hypoxia, regulation of cell death, microglial cell activation (Fig. 5a-b, and Supplementary Table 4). Interestingly, some up-regulated genes only increased in certain cell types. For example, *LIN7A* and *CALD1* were only elevated in cones with aging (Fig. 5c). Additionally, 121 genes related to visual perception, phototransduction, adenosine triphosphate (ATP) biosynthetic process, and retinoid metabolic processes, were down-regulated with aging (Fig. 5d-e, Supplementary Table 4), indicating that visual function is affected as retina ages. We also analyzed the age-related GO change in each cell subtype (Supplementary Fig. 5a-h). Interestingly, in the GO analysis for each cell type, we found most retinal neurons showed similar GOs as total retinal cells (Fig. 5b), such as response to hypoxia, response to oxidative stress and regulation of cell death. In contrast, GOs in MG are anti-apoptosis, negative regulation of programmed cell death and response to hypoxia. These data suggest that retinal cells are heterogeneous during aging. The GO analysis suggested that MG might be in anti-apoptosis status to prevent retinal neuron death during aging. Consistently, among the neurons, a dramatic decrease in rod photoreceptors with retinal aging was observed by scRNA-seq data and immunostaining analysis (Fig. 5f-g), but there was no dramatic cell number change of BCs (Supplementary Fig. 5i-j). Rod outer segments, which are specialized compartment of photoreceptor for phototransduction, were greatly decreased with RHO protein (Fig. 5g). In addition, RHO protein was ectopically expressed in cell bodies, indicating defective light-sensing function of aged rod photoreceptors. In retinal glial cells, the proportion of microglia was remarkably increased in the aged samples (Fig. 5f, h), consistent with the up-regulated genes, which were enriched in response to hypoxia and microglial cell activation (Fig. 5b). Interestingly, the major changes in age-related cell-type proportion in humans were not obvious in the macaque retina (Fig. 5f).

**Figure 5.**
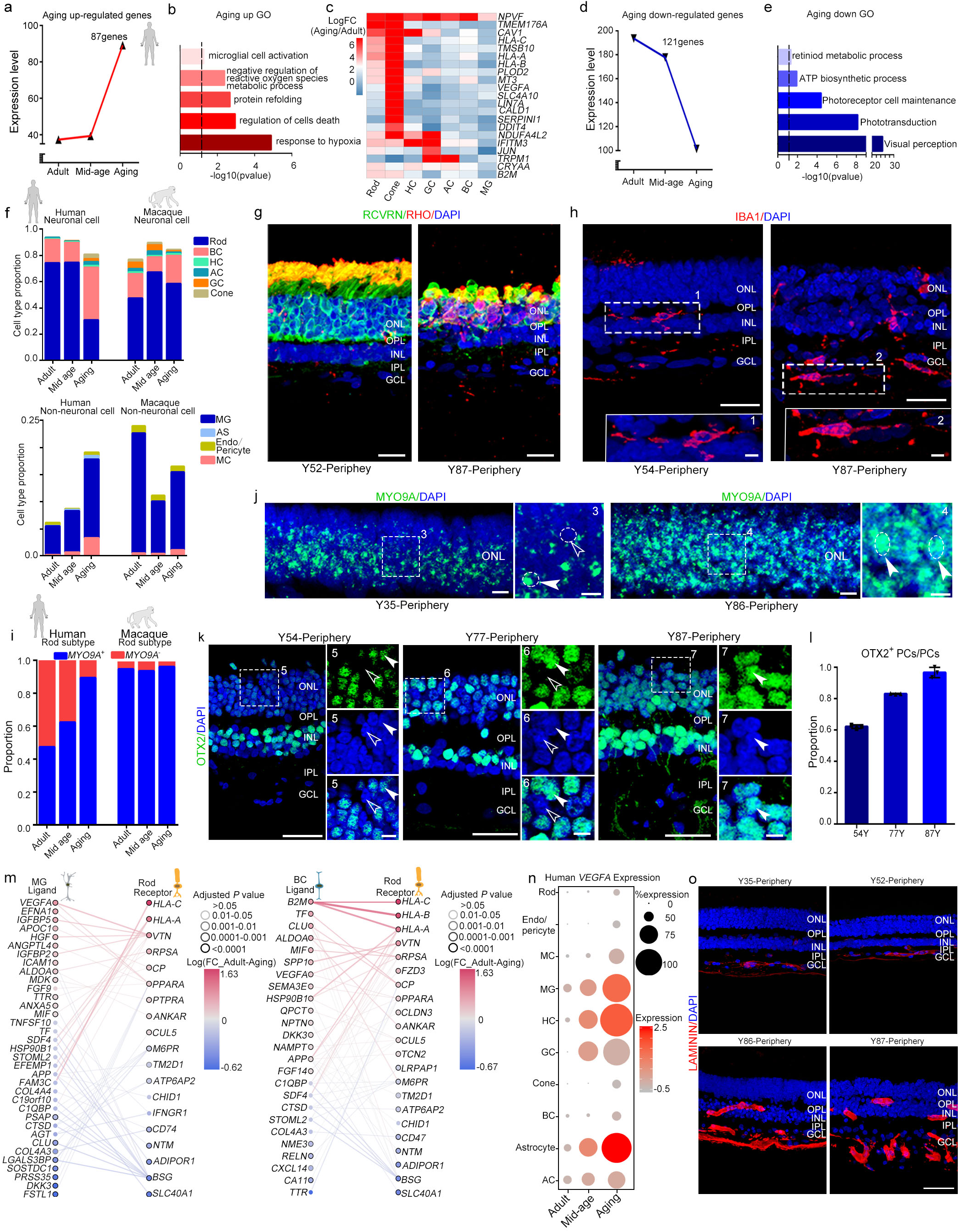
Molecular characteristics of aging primate retina and aging-related changes in cell-cell communication. (a) Co-Expression patterns of up-regulated genes among aging stages. (b) Enriched Gene Ontology terms of up-regulated genes. (c) Heatmap illustrating log2 fold changes of up-regulated genes to cell types at different aging stages in human adult retina. (d) Co-expression patterns of down-regulated genes among aging stages. (e) Enriched Gene Ontology terms of down-regulated genes. (f) Neuronal or non-neuronal cell proportions of human and macaque adult retina at different aging stages. (g-h) Immunostaining of RCVRN/RHO (g) in human retina of Y52 and Y87, IBA1 (h) of Y54 and Y87. Scale bar, 20 μm (g), 25 μm (h, top), 5 μm (h, bottom). Experiments were repeated three times independently with similar results. (i) Proportion of rod subclasses (*MYO9A*^+^ and *MYO9A*^−^) in human adult retina at each stage. (j) *In situ* RNA hybridization of MYO9A in Y35 and Y86 human retina. Blue, DAPI (nucleus marker). Scale bar, 10 μm (left), 5 μm (right). Experiments were repeated three times independently with similar results. (k) Immunostaining of OTX2 in human Y54, Y77, and Y87 peripheral retina. Solid arrowheads indicate positive cells; empty arrowheads indicate negative cells. Scale bar, 25 μm (left), 5 μm (right). Experiments were repeated three times independently with similar results. (l) Quantification of *OTX2^+^* cells in outer nuclear layer (ONL) at various ages, related to Fig. 5k. Data are means ± s.e.m. Each sample was counted from three different slices. (m) Aging-related ligands produced and secreted by MGs with receptors expressed in rods (left) and aging-related ligands produced and secreted by BCs with receptors expressed in rods (right). In all panels, nodes represent ligands or receptors expressed in denoted cell type, and edges represent protein-protein interactions between them. Node color represents magnitude of DEG. Edge color represents sum of scaled differential expression magnitudes from each contributing node, whereas width and transparency are determined by magnitude of scaled differential expression. These figures have been filtered such that top 100 edges representing most differentially expressed node pairs are shown. (n) Dot plot for *VEGF-A* expression in different cell types at each stage in human adult retina. Size of each dot represents percentage of cells in each cluster. Gray to red indicates gradient from low to high gene expression. Experiments were repeated three times independently with similar results. (o) Confocal imaging of LAMININ in human Y35, Y52, Y86, and Y87 peripheral retina. Scale bar, 50 μm. Experiments were repeated three times independently with similar results.

As two major classes of rods were identified in humans (*MYO9A*^+^ and *MYO9A*^−^) (Fig. 2) and rods were the most vulnerable cell type during aging, we next asked whether these two types of rods were differentially affected by aging. Quantification analysis of scRNA-seq data indicated that the population of *MYO9A*^−^ rods dramatically decreased during aging (Fig. 5i), which was confirmed by the ISH of *MYO9A* in the human peripheral retina ONL (Fig. 5j). To further validate *MYO9A*^−^ rods were dramatically decreased, we quantified the ratio of *MYO9A^+^* and *MYO9A*^−^ cells to the total retinal cells in different ages of human retinae (Supplementary Fig. 5k). The data revealed that both *MYO9A^+^* and *MYO9A*^−^ cells were reduced along aging. Moreover, the proportion of *MYO9A*^−^ cells was much more dramatically reduced in aged human retina than *MYO9A^+^* cells. Thus, it appeared that human *MYO9A*^+^ rods were more resistant to aging. Because the macaque rods were predominately *MYO9A*^+^ (95.4%), we did not observe obvious changes in the proportions of *MYO9A*^+^ and *MYO9A*^−^ rods in macaque retina during aging (Fig. 5i). As *OTX2* potentially regulates *MYO9A* expression in human rods (Fig. 2e), we next explored OTX2 expression in the human peripheral retina ONL. Unsurprisingly, *OTX2*^+^ cells in the peripheral ONL increased from 62.1% at 54 years old to 96.7% at 87 years old (Fig. 5k, l). Together, we identified a cell-type-specific degeneration of rod subtypes and depicted gene-expression signatures for each retinal cell type during the process of aging.

### Aging-related changes in cell-cell communication

Aging is a systematic continuous process, regulated by both intrinsic and external signals [43, 44]. To understand how intercellular communication plays a role in the aging of rods, we built a network of ligand-receptor interactions among different cell types and focused on the changes in ligand-receptor expression with aging (Fig. 5m, Supplementary Fig. 5l-m). We studied the ligand-receptor changes between rods and MGs or BCs (Fig. 5m, Supplementary Fig. 5m), as MGs and BCs closely interact with rods. Expression of VEGFA, which are linked to AMD and retinal angiogenesis, were up-regulated in BCs and MGs. Their receptors also increased in rods with aging (Fig. 5m). Interestingly, VEGF-A expression was dramatically increased in MGs, HCs, and astrocytes. Accordingly, abnormal angiogenesis was also observed in the aged retina (Fig. 5n, o, Supplementary Fig. 5n). The length of blood vessels was significantly increased in aged retina. Together, these findings indicate that aging could be a synergistic process with comprehensive changes in different types of neuronal or glial cells in the human retina.

### Molecular changes in foveal and peripheral aged human retina

To investigate the aging trajectories of foveal and peripheral primate retina, we established aging scores using an aging-related gene set which was downloaded from Human Ageing Genomic Resources (https://genomics.senescence.info/). The methods used for aging score calculation was adapted from the algorism used by Nowakowski to predict the maturation of cortical neurons [45] (see Methods). We found that the elevation trends of the aging score generally matched the real sample age (Fig. 6a). Of note, the foveal cells showed higher aging scores than peripheral cells, indicating that the fovea may be more vulnerable to aging. In addition, the increase in aging scores started after 52 years old (Fig. 6a), suggesting that the retinal aging process may gradually accelerate at mid-ages. To verify the reliability of the aging score, we first examined the expression trends of classical aging genes, and found that the expression levels of *STAT3* and *PIK3R1* was positively correlated with aging, whereas the expression levels of *PCNA* and *GSTP1* were negatively correlated with aging, indicating that the aging score models by single-cell transcriptomic profiles were reliable (Fig. 6b, Supplementary Fig. 6a). We then focused on the expression of human aging up-regulated genes in the retina and found that *NPVF, GPX3, B2M, PCP4,* and *CRYAA* were positively correlated with aging, whereas down-regulated genes *RHO, DRD4, GNGT1,* and *MFG8* were negatively correlated with aging (Fig. 6c, Supplementary Fig. 6b-c, Supplementary Table 4).

**Figure 6.**
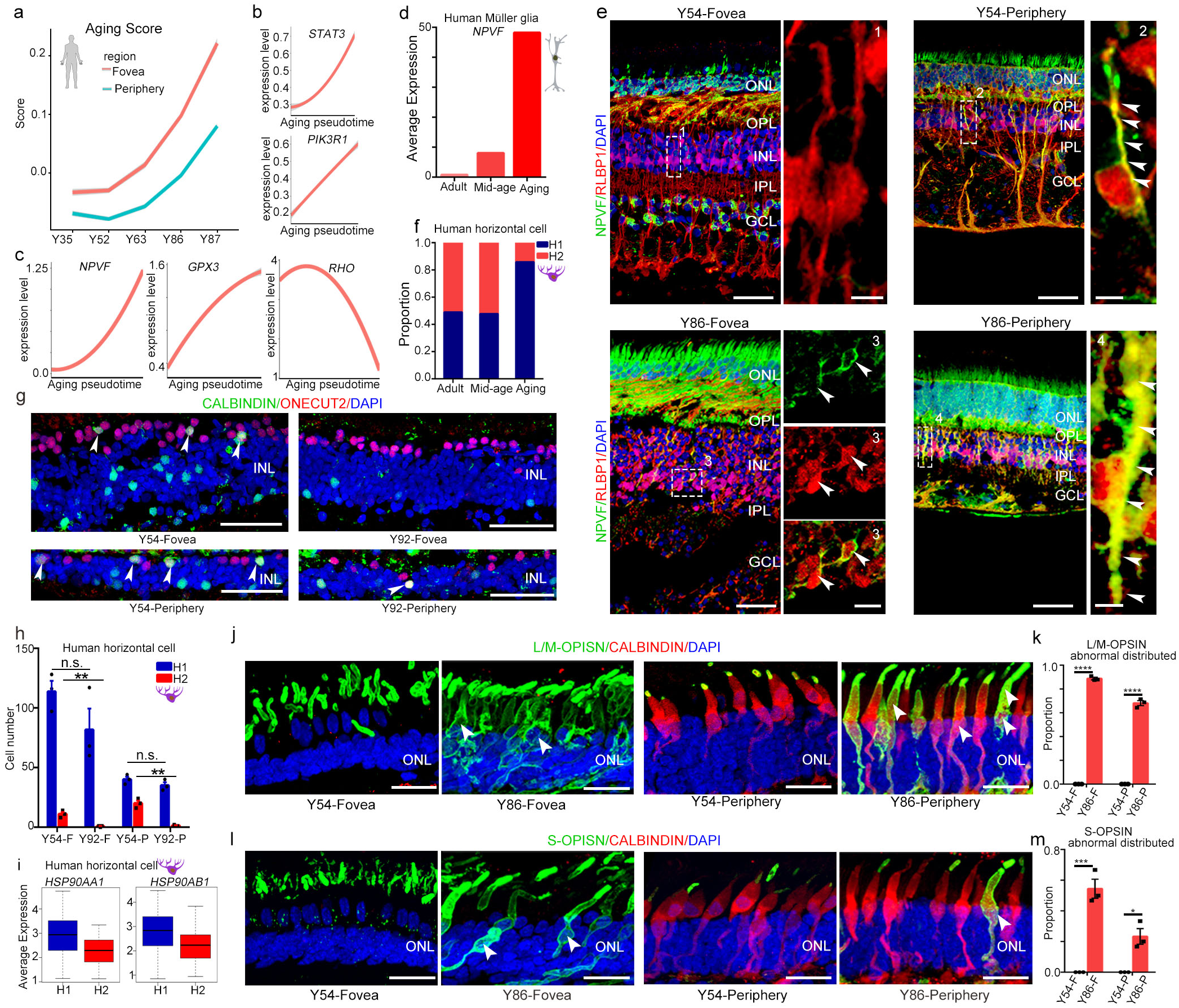
Region- and cell-type-specific molecular changes in aged human retina. (a) Pseudo-aging trajectories between fovea and periphery in human retina. The shadow represents the confidence interval (95%) around the fitted curve. (b) Trajectories of expression of classical human aging genes across human adult retina aging pseudotime. The shadow represents the confidence interval (95%) around the fitted curve. (c) Trajectories of expression of up- and down-regulated genes across human adult retina aging pseudotime. The shadow represents the confidence interval (95%) around the fitted curve. (d) Average expression of *NPVF* at different aging stages in Müller glia. (e) Immunostaining of RLBP1 and NPVF in Y54 and Y86. Solid arrowheads indicate double positive cell. Scale bar of Y54, 50 μm (left), 5 μm (right). Scale bar of Y86, 50 μm (left), 10 μm (right, fovea), 5 μm (right periphery). Experiments were repeated three times independently with similar results. (f) Cell proportions of human H1 and H2 subclasses at different aging stages. (g) Confocal images of adult human retina in Y52 and Y92, Green: CALBINDIN, Red: ONECUT2, Blue: DAPI. Solid arrowheads indicate double positive cell. Scale bar 50 μm. Experiments were repeated three times independently with similar results. (h) Bar chart showing quantification of CALBINDIN/ONECUT2 immunostaining data, related to Fig. 6g. Data are means ± s.e.m, P values calculated by two-sided *t*-test. n.s., no significance, ** *P* <0.01). Each sample was counted from three different slices. (i) Boxplot for *HSP90AA1* and *HSP90AB1* in H1 and H2 subclasses. (j) Immunostaining of L/M-OPSIN/CALBINDIN in Y54 and Y86. Solid arrowheads indicate OPISN translocation in soma. Scale bar, 25 μm. Experiments were repeated three times independently with similar results. (k) Bar chart showing quantification of Fig 6j (data are means ± s.e.m, Y54-F VS Y86-F, **** *P* < 0.0001; Y54-P VS Y86-P, **** *P* < 0.0001). Each sample was counted from three different slices. (l) Immunostaining of S-OPSIN/CABINDIN in Y54 and Y86. Solid arrowheads indicate OPISN translocation in soma. Scale bar, 25 μm. Experiments were repeated three times independently with similar results. (m) Bar chart showing quantification of Fig 6l. Data are means ± s.e.m, P values calculated by two-sided *t*-test. \**P*<0.05, \*\*\**P* <0.001. Each sample was counted from three different slices.

As foveal and peripheral retina exhibited different degrees of aging (Fig. 6a), and MGs, cones, and HCs showed regional differences (see Fig. 3, 4), we investigated whether these regional differences changed during aging. *NPVF*, a highly expressed gene in peripheral MGs, was significantly up-regulated during aging based on the scRNA-seq dataset (Fig. 6d). Consistently, immunostaining of *NPVF* suggested its increased expression in peripheral MGs with aging. Interestingly, *NPVF* was ectopically expressed in the foveal MGs of 86-year-old retina, but not of 54-year-old retina (Fig. 6e). Additionally, we found the average expression of foveal MG-enriched genes gradually decreased during aging, indicating that the regional specificity of foveal MGs declined (Supplementary Fig. 6d). In contrast, the expression of peripheral MG-enriched genes did not show an obvious change trend during aging (Supplementary Fig. 6e).

Other than MGs, the proportion of H2-type HCs in humans decreased from 52.3% (mid-age) to 14.2% (aged) (Fig. 6f). However, this trend was not observed in macaques (Supplementary Fig. 6f), indicating that H2 cells were more sensitive to aging processes in humans than in macaques. We also used immunostaining to verify the changes in HC cell type with aging. The number of H2-type cells decreased in both regions, whereas H1-type cells showed no change (Fig. 6g-h). By comparing the DEGs of H1 and H2 HCs, we found that HSP90AA1 and HSP90AB1, two heat shock proteins, were highly expressed in H1 but not H2 (Fig. 6i). These proteins can mediate lysosomes to refold unfolded proteins, thus protecting cells from the effects of protein toxicity [46]. This may be a reason why the H1 cells showed higher anti-aging ability than H2 cells.

For cones, the regional expression and localization of CALBINDIN showed no differences with aging. However, the subcellular localization of cone OPSINs changed from the outer segments to cell bodies in aged cones (both L/M and S cones) (Fig. 6j-m). It is possible that dysfunction of protein transportation in cilia leads to this mistrafficking and accumulation of cone opsins [47].

### Human retinal disease related to gene expression in retinal cells

We used our scRNA-seq dataset to investigate the cell-type and region specificity of 178 genes associated with 55 types of human retinal disease, including night blindness, macula dystrophy, rod-cone dystrophy, dominant RP, recessive RP, AMD and recessive achromatopsia (RETNET, https://sph.uth.edu/retnet/home.htm). We first calculated the expression of each gene in the foveal or peripheral cell clusters and then aggregated the expression scores by disease types (Fig. 7a-b, Supplementary Fig. 7a-b). Generally, rods and cones, especially foveal cones, were highly related to night blindness and macula dystrophy (Fig. 7a-b). Genes related to rod-cone dystrophy, and dominant and recessive RP were highly expressed in rods and cones, particularly foveal photoreceptors in both humans and macaques. Genes related to AMD were highly enriched in foveal MG and cones, suggesting a correlation of regional cell subtype with this disease.

**Figure 7.**
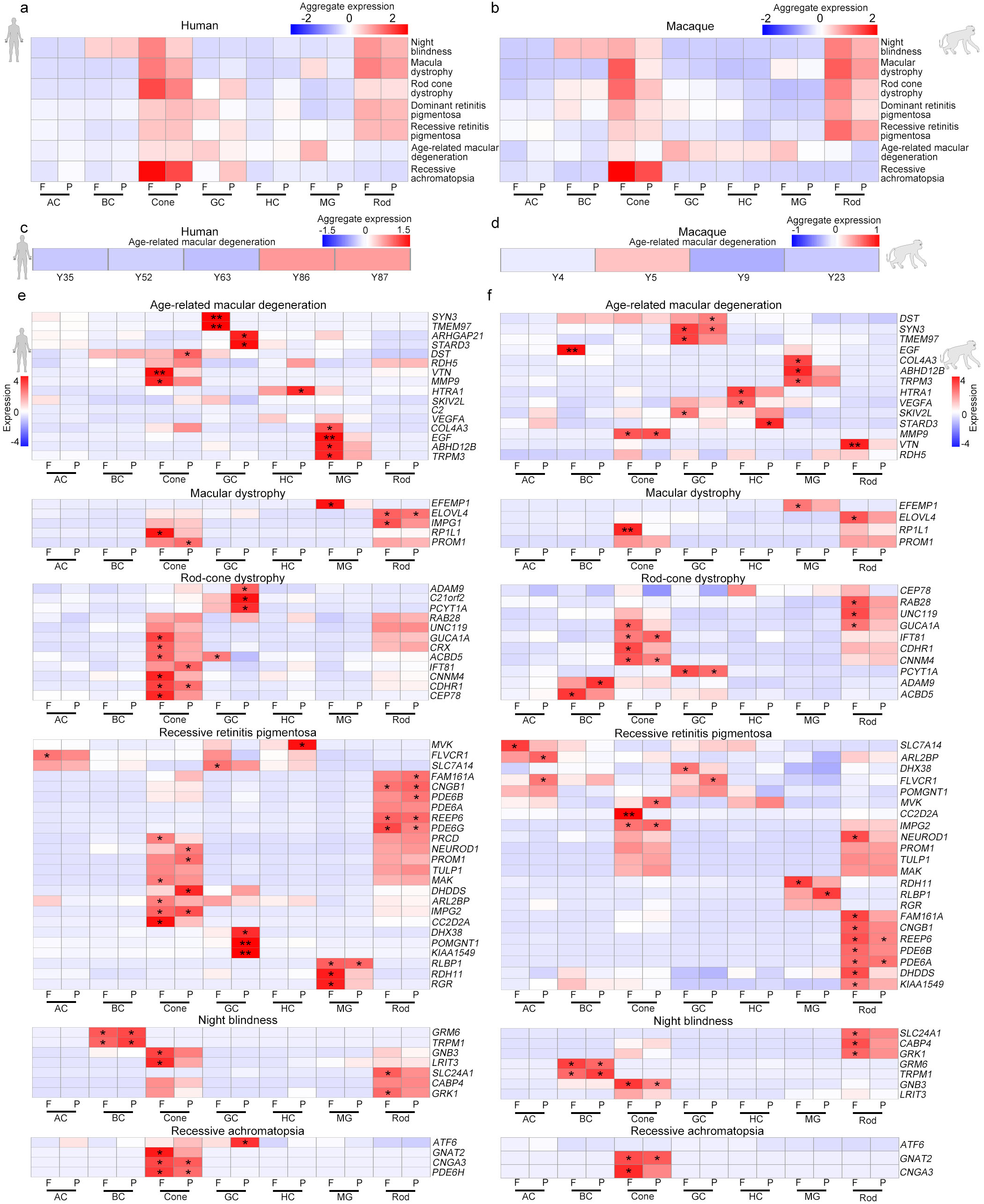
Region-, cell type-, and species-specific expression patterns of genes associated with human retinal diseases. (a-b) Aggregated expression of human eye disease-associated genes in human (a) and macaque (b) cell types, with some diseases showing obvious regional enrichment characteristics. (c-d) Aggregated expression of age-related macular degeneration-associated genes at different human (c) and macaque (d) ages. (e-f) Expression patterns of human eye disease-associated genes in human (e) and macaque (f) retinal cell types. *P* values were calculated by bootstrap hypothesis test, \**P*<0.05, \*\**P*<0.01.

We further illustrated cell-type and region specificity of each gene categorized by disease. As expected, age-related degeneration AMD-risk genes were highly expressed in aged human retina but not in that of aged macaques (Fig. 7c-d), suggesting fewer aging characteristics in the macaque retina. Cell-type and region specificity of disease-related genes were further compared between humans and macaques (Fig. 7e, f, Supplementary Fig. 7a-b). For example, *PDE6A* and *PDE6B*, which are reported to cause recessive RP, were conservatively expressed in the rods in both primates (Fig. 7e-f). *HTRA1* (high-temperature requirement protein A1), which is a risk factor for AMD [48, 49], were highly expressed in HCs in both humans and macaques but with differential regional preference (Fig. 7e-f). Another recessive RP-associated gene, *MVK* (mevalonate kinase), was enriched specifically in human peripheral HCs (Fig. 7e-f). Mutations of *EFEMP1* can lead to macular dystrophy [50], and the expression of *EFEMP1* was high in human and macaque foveal MGs (Fig. 7e-f). In addition, *ATF6*, a recessive achromatopsia-associated gene [51], was selectively expressed in human GCs located in the periphery, but showed no obvious expression in macaques (Fig. 7e-f). Together, these datasets showing region- and cell-type-specific expression patterns of retinal-disease-related genes should be useful for mechanistic and therapeutic studies of retinal diseases in the future.

## Discussion

By profiling 119,520 single cells, we provided detailed single-cell transcriptomes of adult and aged primate retinae with regional information. The retinal transcriptome at single-cell or single-nuclei level has been reported [3, 9–14]. Comparison with other single cell retina studies, we have the largest number of primate retinal samples in one study (6 human samples and 5 macaque samples) covering the longest life span (8 days after birth to 87 years old for humans; 2 - 23 years old for macaques) so far. The non-human primate we used here is *Macaca mulatta*, which is evolutionarily closer to humans than *Macaca fascicularis* [3], adding more primate retinal transcriptome data to the retinal resource. Moreover, we did cross species analysis between humans and macaques, and found the species-specific signatures of primate retinae, which provided valuable data sources to understand the regional and species specializations of the human and macaque retina. In particular, the study provided conceptual advances in the molecular characteristics of aging progress of human retina. Our data revealed that human retinal aging occurred in a region- and cell-type-specific manner, suggesting a strong cell heterogeneity in retinal aging. Moreover, a big dataset was generated and showed the cell-type-, region-, and species-specific gene expression associated with various human retinal diseases, which provides a foundation to understand the molecular and cellular mechanisms underlying human retinal diseases. Overall, this study provides a valuable resource at the single-cell resolution to help understand human and macaque retina and related aging processes, study the regulatory mechanism of the aging process of human retina, and identify molecular markers for aging and degenerative changes of human retina.

### Different rod subtypes in human and macaque retina

We examined retinal cell-type evolutionary conservation between humans and macaques. The major cell types were conserved in the two species, but the molecular features in retinal cells showed species differences (Fig. 1f-k). Based on the expression level of *MYO9A*, the rods in humans and macaques could be divided into two groups. Our data indicated that the *MYO9A*^−^ rod subtype was reduced during aging (Fig. 5i, j), which may result from either depletion *MYO9A*^−^ rods or increased *MYO9A^+^* cells. Considering the proportion of *MYO9A*^−^ cells was much more dramatically reduced in aged human retina than *MYO9A^+^* cells (Fig. 5i, Supplementary Fig. 5k), it is highly possible that the reduction of rods was largely due to depletion of *MYO9A*^−^ rods. Future studies are needed to investigate the functional role of *MYO9A* in rod photoreceptors. It is worth noting that we identified previously unidentified cell types. Cells in Cluster 46 of rod identity and Cluster 51 of MG identity co-expressed *OTX2* and *RLBP1* (Fig. 1d-e, Supplementary Fig. 1e). *OTX2* is a typical marker for BCs and rods, and *RLBP1* is a marker for MGs. MGs are considered as retinal precursor cells in non-mammalian vertebrates [52] and are able to transdifferentiate into rods and BCs with additional transcriptional and epigenetic regulation in mice [53, 54]. The co-expression of *OTX2* and *RLBP1* suggests that MGs in Cluster 51 may be transcriptionally closer to rods and BCs. However, their function and potential to transdifferentiate into neuronal cells, such as rods or BCs, need further exploration.

### Distinct transcriptomes of primate foveal and peripheral retina

With enriched cones, the fovea is the most important region for high-acuity vision in humans, whereas the peripheral retina helps direct eye movements to focus salient images on the fovea [55]. Based on their different functions, they showed distinct molecular signatures in MGs and cones (Fig. 3a, b, h, i). Several genes were found to be specifically enriched in foveal MGs, such as *RGR*, *HTR1*, *DIO2*, and *CYP26A1*. The finding of specific high *RGR* expression in foveal MGs is interesting. *RGR* is a non-visual opsin in intracellular membranes of RPE and MGs [56]. Previous study suggests that RGR opsin and retinol dehydrogenase-10 (*Rdh10*) convert all-trans-retinol to 11-cis-retinol for the regeneration of cone visual pigment in daylight, as RPE cannot meet the higher needs of the visual opsins in rods and cones in daylight [34]. An intriguing speculation is that the higher expression of *RGR* in fovea MGs is initiated to meet the higher demands of visual pigment regeneration in foveal cones.

In addition to *RGR* and retinol, we also found that MGs and cones possibly interact with each other via the TH signaling pathway. As the active form of TH, T3 is important for cone development in retina and retinal organoids [41, 57, 58]. However, it remains unclear whether TH signals exhibit regionally specific distribution in primate retina, and which cell types may provide TH. The major TH in blood is inactive T4, which needs to be converted to T3 by DIO2. Interestingly, we found *DIO2* to be selectively expressed in foveal MGs. Accordingly, we found that cone photoreceptors expressed *THRβ*, a nuclear receptor of T3. Based on our ATAC-seq data, we observed the binding sites of *THRβ* in the promoter regions of *RFC1*, *APLP2*, and *POLE4*, which are genes specifically expressed in foveal cones. Hence, it is possible that the expression of *DIO2* in foveal MGs may produce high local T3, which modulates the expression of a specific set of genes in foveal cones. Of note, higher expression of *NPVF*, which is negatively regulated by the TH signaling pathway [59], was detected in the peripheral cones and MGs. Together, our data imply that foveal MGs may provide a local TH signal that regulates the cellular specificity of foveal and peripheral cones.

### Cellular and molecular changes accompanying human retinal aging

Our study explored cellular and molecular changes during primate retina aging. GO term analysis revealed that microglial cell activation, protein refolding, oxygen species metabolism, and cell death were highly enriched in aging retina and could possibly be considered as hallmarks for retinal aging. In healthy retinae, microglial cells are innate immune cells, which can constantly adapt to their microenvironment. In mouse models, increased density and activation of microglia exist in aged retinae and brains, as well as in age-related human diseases, including AMD, Alzheimer’s disease (AD), and Parkinson’s disease (PD) [60–64]. Similarly, the human retina showed an increased number of microglia during aging (Fig. 5h). Moreover, microglial activation and microglia-mediated inflammatory responses can generate a chronic mild inflammatory environment, including an increased production of inflammatory cytokines, reactive oxygen species (ROS), and reactive nitrogen species (RNS) [65], which can further accelerate retinal aging and pathogenesis of age-related diseases. Thus, we speculated that targeting microglial activation states could potentially be used as a therapeutic avenue in slowing aging and retinal diseases.

Mistrafficking and accumulation of both rod and cone opsins (Fig. 5g and Fig. 6j, l) were found in the aging retina. Damaged protein accumulation can lead to toxicity against photoreceptors as these failed homeostatic mechanisms can contribute to pathology [47]. Therefore, protein misfolding may be a causative reason for apoptosis of photoreceptors during aging. Other than protein misfolding, retinal oxygen supply also decreases with age. Thus, aging retinae suffer from low-grade chronic ischemia [66], which can, in turn, lead to retinal oxidative stress. MGs in the peripheral retina exhibited high expression of metallothioneins (*MT3, MT1G, and MT2A*) and glutathione peroxidase 3 (*GPX3*), which participate in an array of protective stress responses [67]. Metallothioneins are a family of cysteine-rich metal-binding proteins involved in protection against oxidative stress and buffering against toxic heavy metals [68]. As a major scavenger of ROS in plasma, *GPX3* acts as a redox signal modulator [69]. The high levels of metallothioneins and *GPX3* in the peripheral MGs may contribute to fovea to peripheral aging gradient observed in human retina.

VEGF is a well-known risk factor for abnormal retinal angiogenesis in AMD and diabetic retinopathy. We found that VEGF-A expression increased specifically in HCs, MGs, and astrocytes following aging. VEGF is a potent endothelial cell mitogen that stimulates proliferation, migration, and tube formation, leading to angiogenic growth of blood vessels [70]. With the knowledge that cellular origin of VEGF-A increases during aging, it is possible to design therapeutic approaches to reduce its expression in MGs, HCs, and astrocytes for abnormal retinal angiogenesis treatment. Other than VEGF, we observed several previously unidentified genes associated with aging, such as *NPVF*, *PCP4*, *CRYAA*, and *B2M*. Together, our study provides a valuable data source for future studies on human retinal aging and related diseases.

### Cell-type and region specificity of human retinal diseases

Our study provides a foundation to understand the molecular and cellular mechanisms underlying human retinal diseases. By assessing the expression of 178 genes implicated in human retinal diseases, we found that these genes were preferentially expressed in specific retinal cell classes and regions. Photoreceptors in foveal regions are the major cell types for many retinal degenerative diseases. These findings emphasized the value of human aging transcriptomes for age-related retinal diseases. In particular, AMD-risk genes were highly expressed in the aged human retina. Together, the region- and cell-type-specific expression patterns of retinal disease-related genes may provide guidelines for future cellular and molecular precision therapies.

## Acknowledgments

We thank the National Human Brain Bank for Development of Function. This work was supported by the Strategic Priority Research Program of the Chinese Academy of Sciences (XDA16020601/03), National Key R&D Program of China (2019YFA0110100, 2017YFA0103303, 2017YFA0102601), National Natural Science Foundation of China (NSFC) (81925009, 81790644, 31671072, 81891001, 81900855, 31771140), Fundamental Research Funds for the Central Universities (WK2070000174), and Beijing Brain Initiative of Beijing Municipal Science & Technology Commission (Z181100001518004).

## Author contributions

T.X., X.W., Q.W., and M.Z. conceived the project and designed the experiments. W.Y., S.Z., M.W., H.X., and M.Z. performed RNA-seq. Y.L., H.D., and M.W. analyzed the data. W.Y. and Q.W. performed immunofluorescence and imaging. W.Y. and H.X. prepared the ATAC-seq library. W.Y. and Y.L. performed RNA-scope ISH. Z.Y., Y.T., H.Q., R.P., J.H., Y.M., S.L., W.W., Q.M., Z.L., C.Y., S.L., J.W., and C.M. prepared the primate samples. T.X., X.W., Q.W., and M.Z. wrote the manuscript with input from all other authors. All authors edited and proofread the manuscript.

## Methods

### Human retinal samples

The isolation procedure of human retinal samples and research protocols were approved by the Research Ethics committee of the Peking University Third Hospital and Peking Union Medical College Hospital, and were conducted in accordance with approved institutional guidelines. All the protocols are in compliance with the “Interim Measures for the Administration of Human Genetic Resources”, administered by the Ministry of Science and Technology of China.

### Macaque retinal samples

Rhesus macaques (*Macaca mulatta*) were provided by Xieerxin Biology Resource (Beijing, China). All procedures were conducted in accordance with the Principles for the Ethical Treatment of Non-Human Primates and was approved by the Institutional Animal Care and Use Committee of the Institute of Biophysics and University of Science and Technology of China. All animals were maintained at 25°C on a 12 h light:12 h dark schedule at Xieerxin Biology Resource, a Laboratory Animal Care-accredited facility in Beijing, in compliance with all local and federal laws governing animal research. Animals were given a commercial diet twice a day with tap water provided *ad libitum* and were fed vegetables and fruits once daily under careful veterinary oversight. Before the experiment, none of the animals had a clinical or experimental history that would affect physiological aging or increase disease susceptibility.

### Tissue sample collection and dissociation

Human and macaque eye tissue samples were collected in ice-cold artificial cerebrospinal fluid (ACSF) containing 125.0 mM NaCl, 26.0 mM NaHCO_3_, 2.5 mM KCl, 2.0 mM CaCl_2_, 1.0 mM MgCl_2_, 1.25 mM NaH_2_PO_4_; pH 7.4, bubbled with carbogen (95% O_2_ and 5% CO_2_). The sclera, RPE, lens, and vitreous body were then removed. After separation of the whole retina, the retinal fovea (macula) (1.5–2.0 mm) was identified by full of lutein. The peripheral and foveal retina were gently separated into small pieces and centrifuged at 200 g for 2 min. The supernatant was removed, to which was added 500 μl of digestion buffer (2 mg/ml collagenase IV (Gibco), 10 U/μl DNase I (NEB), and 1 mg/ml papain (Sigma) in phosphate-buffered saline (PBS)). The tissue samples were rotated and incubated at 37 °C on a thermo-cycler at 300 g for 20–25 min. The sample was pipetted every 5 min [71] to digest the tissue into single cells.

### Library preparation for high-throughput sequencing

Cells were suspended in 0.04% bovine serum albumin (BSA)/PBS at the proper concentration to generate cDNA libraries with Single Cell 3’ Reagent Kits, according to the manufacturer’s instructions. Thousands of cells were partitioned into nanoliter-scale Gel Bead-In-EMulsions (GEMs) by 10x™ GemCode™ Technology, in which the cDNA produced from the same cell shared a common 10x Barcode. Upon dissolution of the Single Cell 3’ Gel Bead in GEM, primers containing an Illumina R1 sequence (read 1 sequencing primer), 16-bp 10x Barcode, 10-bp randomer, and poly-dT primer sequence were released and mixed with cell lysate and Master Mix. After GEM incubation, barcoded, full-length cDNA from poly-adenylated mRNA was generated. The GEMs were then broken, and silane magnetic beads were used to remove leftover biochemical reagents and primers. Prior to library construction, enzymatic fragmentation and size selection were used to optimize the cDNA amplicon size. P5, P7, an index sample, and R2 (read 2 primer sequence) were added to each selected cDNA during end repair and adaptor ligation. The P5 and P7 primers were used for Illumina bridge amplification of the cDNA (http://10xgenomics.com). Finally, the library was processed on the Illumina HiSeq4000 platform for sequencing with 150-bp pair-end reads.

### Single-cell RNA-seq data preprocessing

Cell Ranger v2.0.1 (http://10xgenomics.com) was used to process the raw sequencing data with default parameters. Human reads were aligned to the human reference genome (hg19). We used Cell Ranger to create a pre-mRNA reference, using an available transcriptome from the Ensembl genome browser for *M*. *mulatta* (annotation release 95). We excluded poor quality cells after the gene-cell data matrix was generated by Cell Ranger with Seurat (v2.2.0) (https://satijalab.org/seurat/pbmc3k_tutorial.html) in Bioconductor [72, 73]. Only cells that expressed more than 500 genes and fewer than 6 000 genes were considered, and only genes expressed in at least 0.01% of total cells were included for further analysis. Cells with mitochondrial gene percentages over 30% were discarded as well. In total, 22 347 genes across 38 558 human single cells and 18 933 genes across 80 962 macaque single cells remained for subsequent analysis. The data were natural log-transformed and normalized to a total of 1e4 molecules per cell for scaling sequencing depth using Seurat. Batch effects were mitigated using the ScaleData function in Seurat.

### Identification of cell types and subtypes by dimensional reduction

Seurat (v2.2.0) was used to perform linear dimensional reduction. Highly variable genes with average expression between 0.0125 and 8 and dispersion greater than 1 were selected as inputs for principal component analysis (PCA). We then determined the significant PCs using the JackStrawPlot function. The top 20 PCs were applied for t-Distributed Stochastic Neighbor Embedding (tSNE) to cluster cells with the FindClusters function in Resolution 4.0. Clusters were identified by the expression of known cell-type markers.

### Identification of differentially expressed genes (DEGs) among clusters

DEG analysis among clusters was performed with the Seurat function FindAllMarkers (thresh.use = 0.25, test.use = “bimod”). The bimod [74] test returns likelihood-ratios for single-cell gene expression, and genes with an average expression difference >0.25 natural log and *P* < 0.05 were selected as marker genes. Enriched Gene Ontology (GO) terms of marker genes were identified using DAVID v6.7 [75, 76] (https://david.ncifcrf.gov/home.jsp).

### LIGER analysis for human and macaque datasets

To evaluate the conservation and variation between the human and macaque retina transcriptomes, we used LIGER [25] (https://macoskolab.github.io/liger/) to integrate human and macaque retina datasets with the function createLiger. The function selectGenes (var.thresh = 0.1) was used to perform variable gene selection on human and macaque datasets separately and then in combination. We next identified cells loaded on corresponding cell factors and quantile-normalized their factor loadings across datasets. Cell dimensionality reduction was performed with the function runTSNE. The function plotGeneLoadings was used to visualize the most highly loaded genes (both shared and dataset-specific) for each factor. To compare different cluster assignments, we employed the function makeriverplot to visualize the previous cell-type assignments of humans and macaques with liger joint clusters.

### Gene expression correlation analysis between human and macaque cell types

To explore the similarities between human and macaque cell types, we calculated the Pearson correlation coefficients across humans and macaques with shared variable genes in both datasets. The resulting correlation matrices were visualized with circus plot using the R Package circlize.

### WGCNA in regional DEGs of human MGs

To identify the human MG regionally related gene modules, we identified the DEGs between foveal and peripheral MG, and then obtained various gene modules under WGCNA [77, 78] (https://cran.r-project.org/src/contrib/Archive/WGCNA). Module assignment was followed by quantifying the relationship between modules and region traits, where the correlations among them were computed and shown as a heatmap. The blue module had the closest association with the fovea. The blue module network was plotted using Cytoscape.

### Co-expression analysis for aging-related genes

We divided the samples into adult (Y35, Y52, Y63) and aged groups (Y86, Y87). We then performed FindAllMarkers using the Seurat package to identify DEGs. Only genes with an average expression difference greater than 0.5 were selected as aging-related genes. The up-regulated (enriched in aged group) and down-regulated (enriched in adult group) age-related genes were calculated by mean expression among different aging stages, respectively.

### Intercellular network analysis

Cell-cell interactions were predicted using a method similar to that described previously [22, 79]. We created a cell communication interactome and collected known protein-protein interactions between receptor and ligand and all related genes were collected [22]. Gene lists were manually filtered with the DEGs of the adult to aged groups in our cell types. To investigate aging-related perturbations in the putative cell-cell interaction networks, DEG metrics from the MAST analysis outlined above were used to build subnetworks for each set of interactions between cell types. In these networks, nodes represent ligands or receptors expressed in the denoted cell type, and edges represent protein-protein interactions between them. Nodes were colored to represent the magnitude of DEGs. These values were scaled per cell type and summed to determine edge weight.

### Assignment of pseudo-aging score for human retina cells

Aging-related genes were downloaded from Human Ageing Genomic Resources (https://genomics.senescence.info/). We performed PCA with aging-related genes using the function RunPCA and determined statistically significant PCs using the function JackStraw. We then computed the correlation between the age vector and significant PCs and then selected the PC resulting in the highest correlation coefficient. Pseudo-aging scores of cells were determined by the average expression of the chosen PC genes.

### Analysis of cell-type specific expression of diseases

The human retina disease genes were obtained from the Retinal Information Network (http://www.sph.uth.tmc.edu/RetNet/). For each of the foveal and peripheral cell types that belonged to humans and macaques, we first calculated the expression of all genes across all cell types and computed the fraction of cells in each cell type. We then obtained a matrix of gene expression scores for all genes across all cell types, and selected retinal disease related genes to visualize expression patterns. We computed the mean relative expression strengths among each cell type for the different retinal diseases (e.g., night blindness, macula dystrophy, rod cone dystrophy, dominant RP, recessive RP, AMD, recessive achromatopsia). The calculation method was described by Peng et al.[3]. Finally, we calculated *P* values across different cell types for each disease gene by using bootstrap hypothesis test.

### ATAC library preparation for high-throughput sequencing

ATAC-seq was performed as described previously [80, 81]. In total, 50,000– 60,000 cells were twice washed with 50 μl of cold PBS and immediately resuspended in 50 μl of ATAC-lysis buffer (10 mM Tris-HCl pH 7.4, 10 mM NaCl, 3 mM MgCl_2_, 0.1% (v/v) Nonidet P40 Substitute) and centrifuged for 10 min at 500 g at 4 °C. The resulting supernatant was then removed and added to 50 μl of transposition reaction mix (10 μl 5 × TTBL buffer, 4 μl TTE Mix, and 36 μl nuclease-free H_2_O) of a TruePrep DNA Library Prep Kit V2 (Vazyme TD501-02). Samples were then incubated at 37 °C for 30 min. After the reaction finished, deoxynucleotide was isolated using a QIAquick PCR Purification Kit (QIAGEN 28106). The ATAC-seq libraries were then prepared using the Trueprep DNA Library Prep kit V2 (Vazyme TD501-02). Finally, 2 × 150 paired-end sequencing was performed on an Illumina HiSeq X-10.

### ATAC-seq data analysis

We cleaned the reads using fastp (v0.19.6) and then mapped the reads to the hg19 reference genome with the parameters: -t -q -N 1 -L 25 -X 2000 using Bowtie2 (v2.3.4.3). All unmapped reads, non-uniquely mapped reads, and polymerase chain reaction (PCR) duplicates were removed. Fragments mapped to blacklisted genomic regions were removed. The uniquely mapped reads were shifted +4/−5 bp according to the strand of the read. To visualize the ATAC-seq signal, we extended each read by 50 bp and counted the coverage for each base. All ATAC-seq peaks were called by MACS2 v2.1.1.

### ATAC-seq data quality control

Quality of the ATAC-seq data was evaluated for several parameters, including number of raw reads, alignment rate, percentage of reads mapped to chromosome M, percentage of reads mapped to repeat regions (black list), percentage of reads passed MAPQ score filter, percentage of total signal within known artefact regions, and correlation between replications.

### Connecting transcription factors to target genes

To find the potential transcription factors binding to *MYO9A*, the *THRB* regulatory sequence, FIMO from MEME Suite (v5.0.5) was used for motif enrichment analysis with default parameters.

### Immunohistochemistry

Human retinal tissue samples were fixed overnight in 4% paraformaldehyde. The fixed retinae were dehydrated in 20% and 30% sucrose in PBS at 4 °C and embedded in optimal cutting temperature medium (Thermo Scientific). Thin 20– 25 μm cryosections were collected on superfrost slides (VWR) using a Leica CM3050S cryostat. For immunohistochemistry, antibodies against the following proteins were used at the indicated dilutions: Mouse anti-RLBP1 (1:500, Abcam), Mouse anti-Rod-OPSIN (1:1 000, Sigma), Rabbit anti-S-OPSIN (1:500, Millipore), Rabbit anti-L/M-OPSIN (1:500, Millipore), Mouse anti-Calbindin (1:500, Abcam), Rabbit anti RRKCA (1:500, Abcam), Sheep anti VSX2 (1:400, Exalpha Biologicals), Sheep anti ONECUT2 (1:40, R&D Systems), Goat anti OTX2 (1:200 R and D Systems), Rabbit anti-RCVRN (1:500, Millipore), Rabbit anti-NPVF (1:500, Sigma), Rabbit anti-TRH (1:500, Sigma), Rabbit anti BTG1 (1:100, SAB biotech), Mouse anti NR2E3 (1:100, R&D Systems) and Goat anti IBA1 (1:500 Abcam). Primary antibodies were diluted in blocking buffer containing 10% donkey serum, 0.2% Triton X-100, and 0.2% gelatin in PBS at pH 7.4. Alexa Fluor 488, Alexa Fluor 594, or Alexa Fluor 647 fluorophore-conjugated secondary antibodies (1:500) (Life Technologies) were used as appropriate. Cell nuclei were stained with DAPI (1:10 000). Images were collected using an Olympus FV1000 and Olympus FV3000 confocal microscope (Japan).

### RNAscope

RNAscope® detection was conducted in strict accordance with the ACD RNAscope® protocols [82]. Fresh retinal sections were dehydrated in sequential incubations with ethanol, then repaired in boiling repair solution for 5 min, followed by 30 min protease III treatment and washing in ddH_2_O. Appropriate combinations of hybridization probes (CYP26A1 Cat#487741; MYO9A Cat#518511; RLBP1 Cat#414221-C2; DIO2 Cat#562211-C3; LHX1 Cat#493021; ISL1 Cat#478591-C2)were incubated for 2 h at 40 °C, followed by fluorescence labeling, DAPI counterstaining, and mounting with Prolong Gold mounting medium.

### Statistical analyses

All data were represented as means ± s.e.m. Comparisons between two groups were made using *t*-tests. Quantification graphs were analyzed using GraphPad Prism (GraphPad Software).

### Data and Code availability

The single-cell RNA-seq and bulk ATAC-seq data used in this study were all deposited in the Genome Sequence Archive (GSA) for Human, National Genomics Data Center (https://bigd.big.ac.cn/gsa-human) under BioProject accession numbers PRJCA002731. GSA number for macaque is CRA002680 and GSA number for Human is HRA000182. All codes for data analysis are included in Supplementary file 1.

---

**Supplementary Figure 1.**
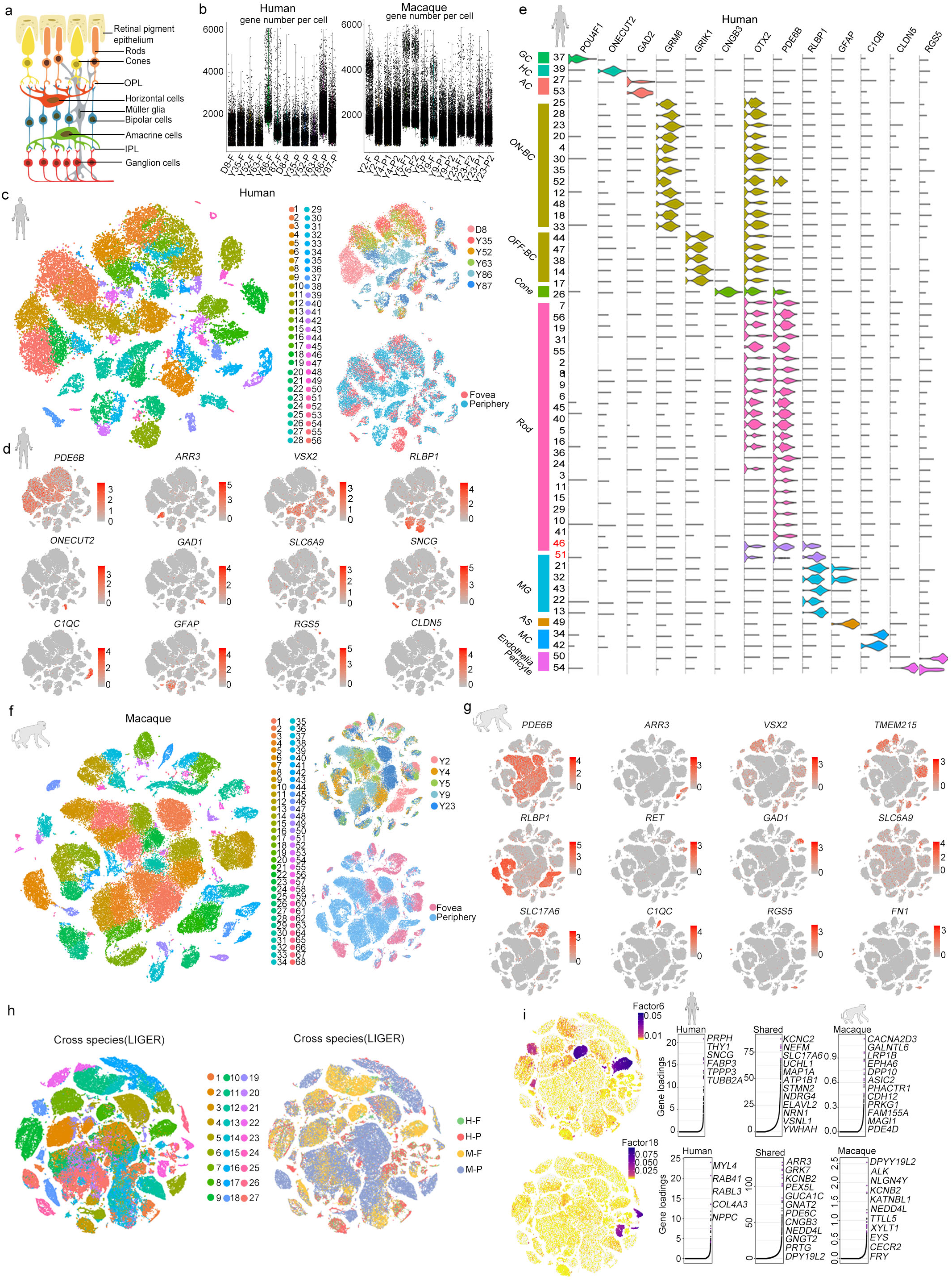
Single-cell RNA-seq information and molecular diversity of primate retina. (a) Schematic of major cell types and three-layer construction in retina. (b) Quality control for human and macaque samples, with each dot representing a single cell (F: Fovea; P: Periphery). (c) t-SNE visualization of human retina samples colored by cluster (left), age, and region (right). Each dot represents an individual cell. (d) Expression patterns of known markers for different cell types in human adult retina displayed in t-SNE plots (gray, no expression; red, relative expression). (e) Violin plots showing expression of different cell-type marker genes, distinguishing 56 subclasses in human adult retina. (f) t-SNE visualization of macaque retina samples colored by cluster (left), age, and region (right). Each dot represents an individual cell. (g) Expression patterns of known markers for different cell types in macaque adult retina displayed in t-SNE plots. (h) t-SNE visualization of 119 520 single cells analyzed by LIGER, color-coded by LIGER joint clusters (left) and species and regions (right) (H-F: human-fovea; H-P: human-periphery; M-F: macaque-fovea; M-P: macaque-periphery). (i) Cell factor loading values (left) and gene loading plots (right) of LIGER joint clusters and shared or species-specific genes for factor 6 and factor 18.

**Supplementary Figure 2.**
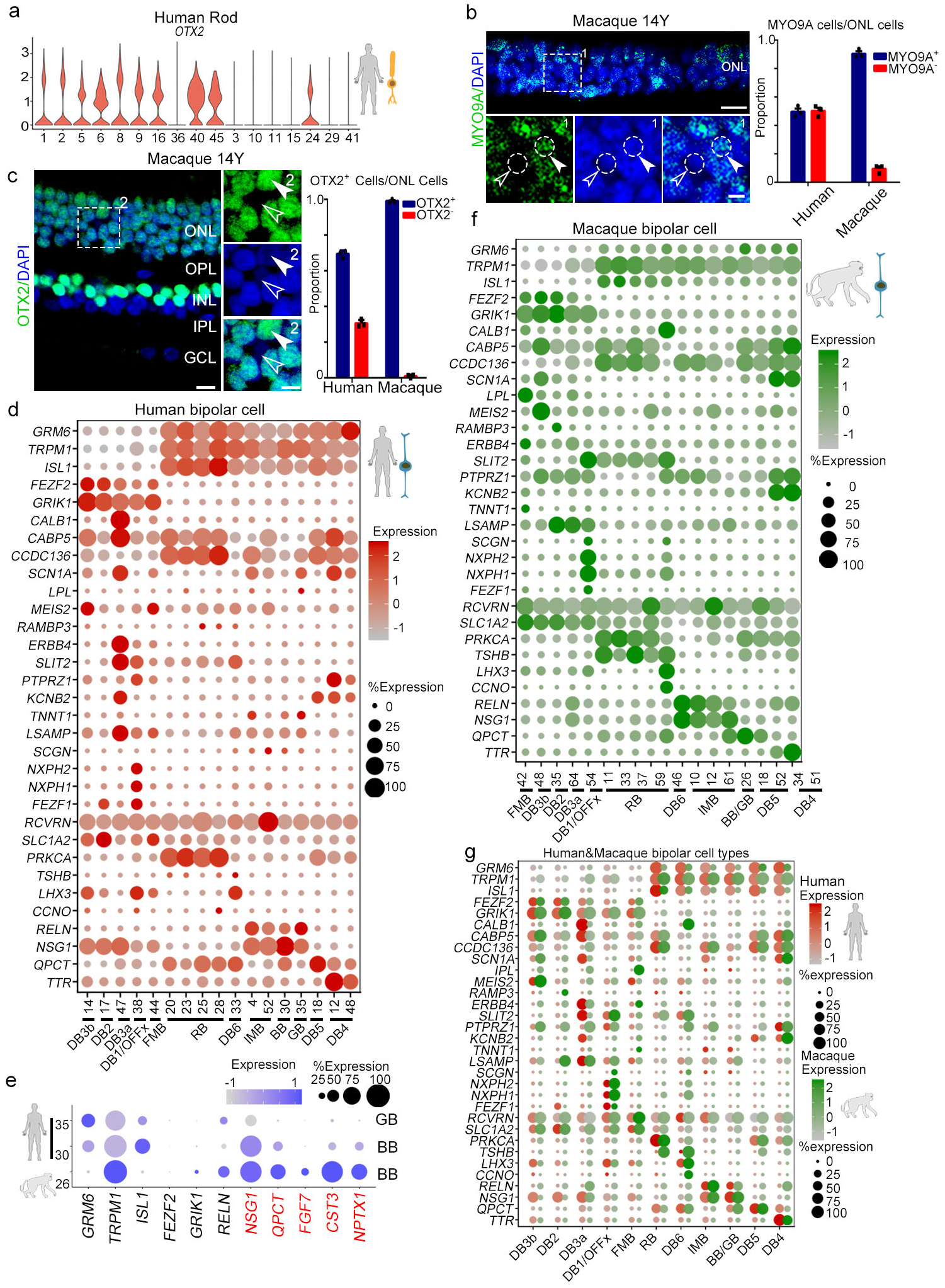
Distinct subtypes of human and macaque bipolar cells. (a) Violin plots showing expression of marker gene *OTX2*, distinguishing 17 rod subclasses in human adult retina. (b) Left: confocal imaging of *MYO9A* in adult macaque retina. Solid arrowheads indicate *MYO9A^+^* cells; empty arrowheads indicate *MYO9A*^−^ cells. Blue, DAPI (nucleus marker). Scale bar, 10 μm (top), 5 μm (bottom). Experiments were repeated three times independently with similar results. Right: Bar chart showing the quantification of Fig. 2d and supplementary Fig. 2b. Data are means ± s.e.m. Each sample was counted from three different slices. (c) Left: confocal imaging of OTX2 in adult macaque retina. Solid arrowheads indicate OTX2^+^ cells; empty arrowheads indicate OTX2^−^ cells. Blue, DAPI (nucleus marker). Scale bar, 10 μm (left), 5 μm (right). Experiments were repeated three times independently with similar results. Right: Bar chart showing the quantification of Fig. 2f (Y-52 periphery) and supplementary Fig. 2c. Data are means ± s.e.m. Each sample was counted from three different slices. (d) Dot plot for broad markers and type-enriched markers of different bipolar cell clusters in human adult retina. Color of each dot shows average scale expression, and its size represents percentage of cells in cluster. (e) Dot plot for type-enriched markers of BB/GB subclasses in human and macaque adult retina. Color of each dot shows average scale expression, and its size represents percentage of cells in cluster. (f) Dot plot for broad markers and type-enriched markers of different bipolar cell clusters in macaque adult retina. Color of each dot shows average scale expression, and its size represents percentage of cells in cluster. (g) Dot plot for broad markers and type-enriched markers of different bipolar cell subclasses in human (red) and macaque (green) adult retina. Size of each dot represents percentage of cells in each cluster. Gray to red/green indicates gradient from low to high gene expression.

**Supplementary Figure 3.**
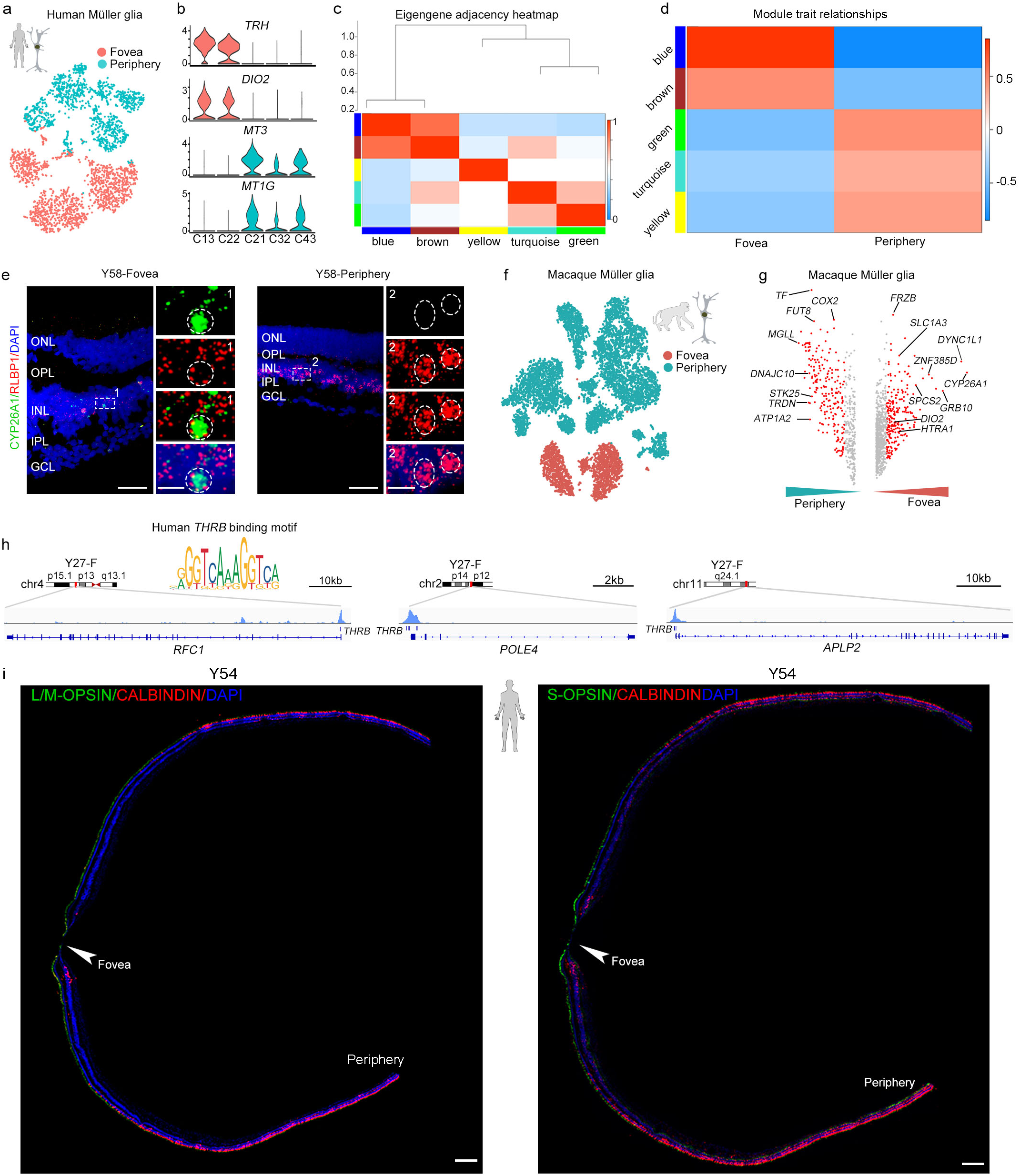
Molecular properties of primate MGs and cones. (a) t-SNE plot of human Müller glia distinguished by regions (dots, single cell; color, regions). (b) Violin plots showing expression of DEGs that distinguish Müller glia subclasses in human adult retina. (c) Gene dendrogram obtained by average linkage hierarchical clustering. Color underneath row corresponds to module assignment; each color represents different assigned module. (d) Module-trait associations. Each row corresponds to a module eigengene, column to a trait. Each cell contains corresponding correlations and *P* values. Color key from blue to red indicates low to high correlation. (e) *In situ* RNA hybridization of *CYP26A1* and *RLBP1* in Y58 human retina. Blue, DAPI (nucleus marker). Scale bar, 50 μm (left), 10 μm (right). (f) t-SNE plot of macaque Müller glia distinguished by regions (dots, single cell; color, regions). (g) Volcano plot for DEGs of macaque foveal and peripheral Müller glia. Red dots show average log2 fold changes >0.5. (h) Normalized ATAC-seq profiles of *RFC1*, *POLE4*, and *APLP2* in Y27 foveal retina showing activation of these genes. DNA binding motif of *THRB* (top), identified in ATAC-seq peaks close to *THRB* transcription start site (TTS). (i) Immunostaining of L/M-OPSIN/CALBINDIN and S-OPSIN/CABINDIN in Y54, Scale bar, 300 μm. Experiments were repeated three times independently with similar results.

**Supplementary Figure 4.**
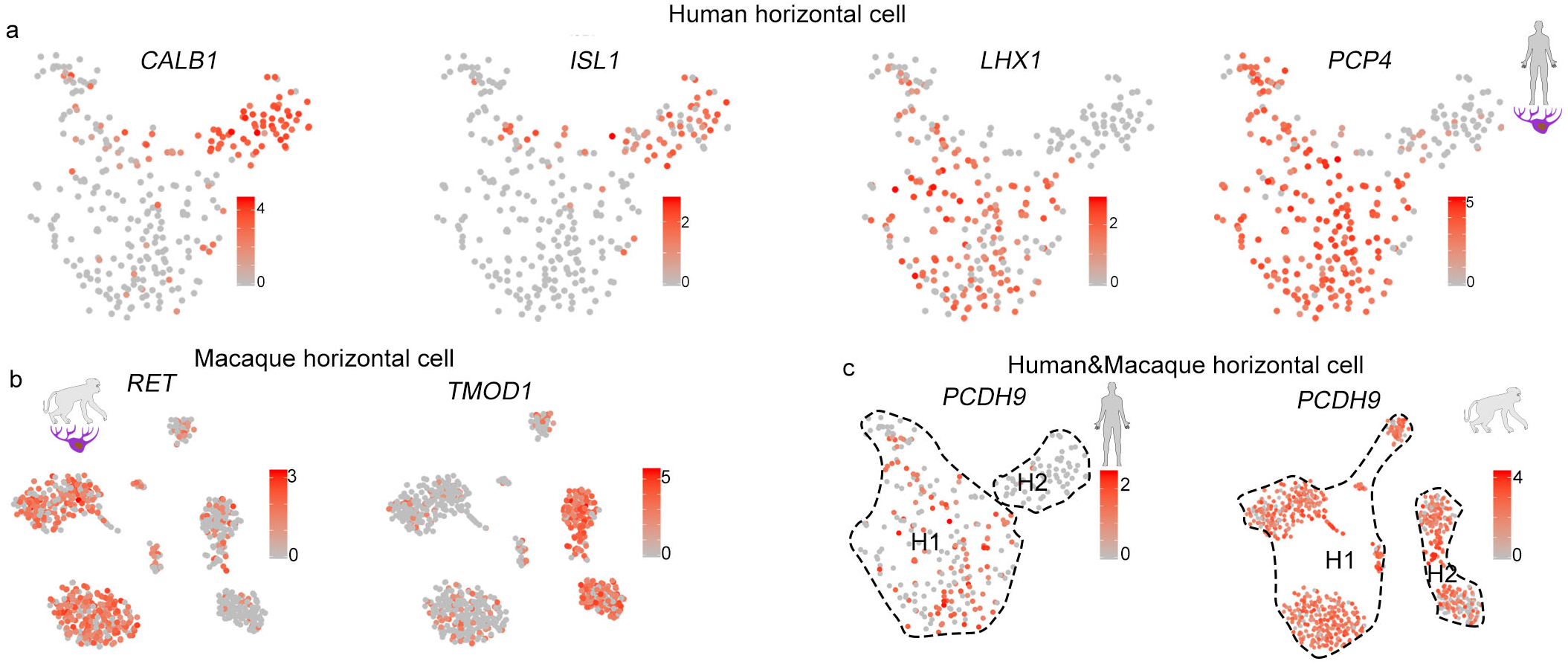
Single-cell profiles of retinal horizontal cells in primates. (a) Visualization of expression of known marker genes for horizontal cells by t-SNE in human adult retina. Cells are colored according to gene expression levels. (b) Visualization of expression of known marker genes for horizontal cells by t-SNE in macaque retina. Cells are colored according to gene expression levels. (c) Visualization of expression of *PCDH9* for human (left) and macaque (right) horizontal cells by t-SNE. Cells are colored according to gene expression levels.

**Supplementary Figure 5.**
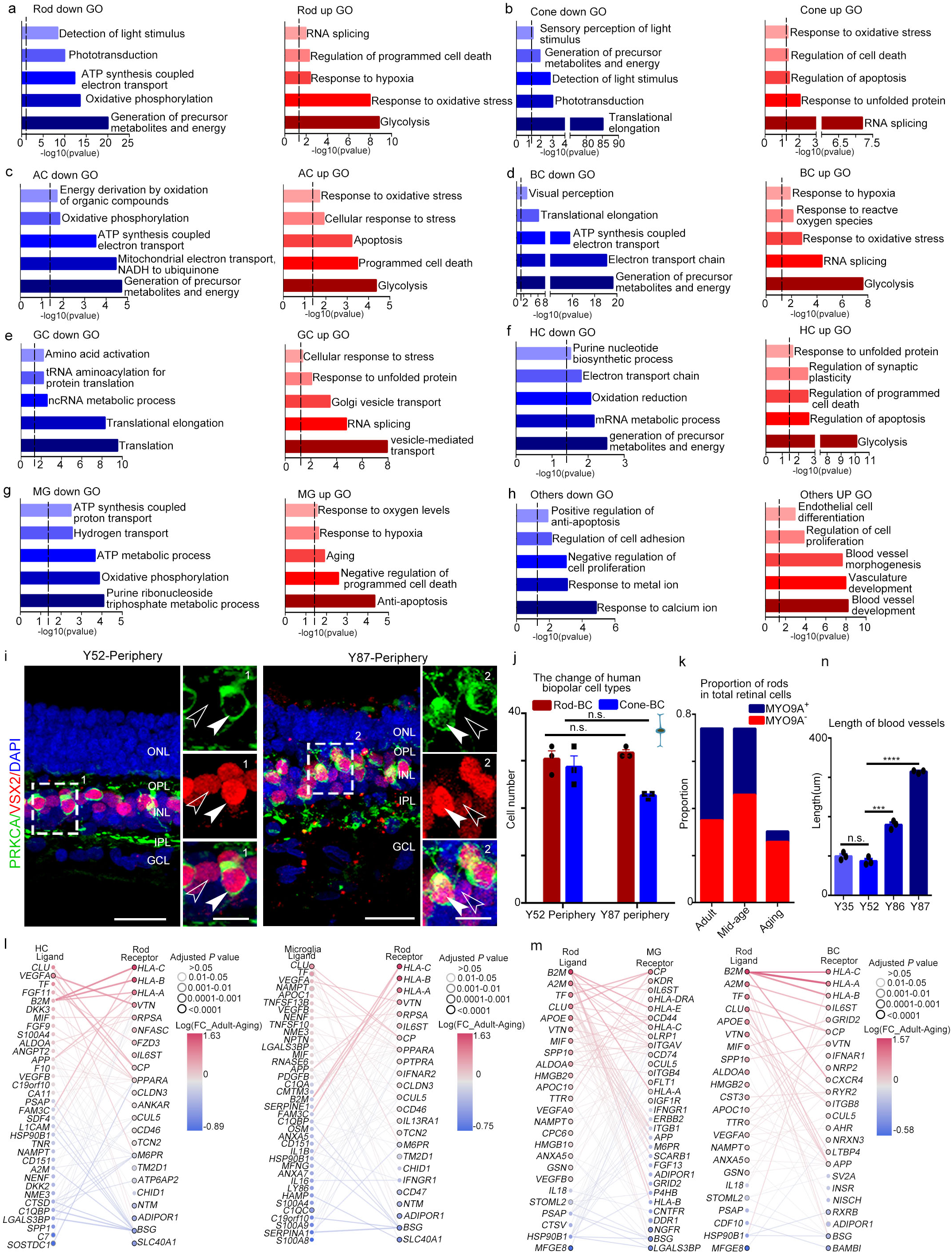
Cell-type-specific aging patterns in human retina and aging-related changes in cell-cell communications. (a-h) Enriched GO terms of cell type specific aging up/down-regulated genes. a(Rod), b(Cone), c(AC), d(BC), e(GC), f(HC), g(MG), h(Others). (i) Immunostaining of PRKCA and VSX2 in human Y52 and Y87 peripheral retina. Solid arrowheads indicate double positive cells; empty arrowheads indicate PRKCA negative cells. Scale bar, 25 μm (left), 10 μm (right). Experiments were repeated three times independently with similar results. (j) Bar chart showing quantification of Supplementary Fig.5i. Data are means ± s.e.m, P values calculated by two-sided *t*-test. n.s., no significance. Each sample was counted from three different slices. (k) Proportion of rod subclasses (*MYO9A*^+^ and *MYO9A*^−^) in total human retinal cells at each stage. (l) Aging-related ligands produced and secreted by rods with receptors expressed in MGs (left) and aging-related ligands produced and secreted by rod with receptors expressed in BCs (right). (m) Aging-related ligands produced and secreted by HCs with receptors expressed in rods (left) and aging-related ligands produced and secreted by microglia with receptors expressed in rods (right). In (l) and (m) panels, nodes represent ligands or receptors expressed in denoted cell type, and edges represent protein-protein interactions between them. Node color represents magnitude of DEGs. Edge color represents sum of scaled differential expression magnitudes from each contributing node, whereas width and transparency are determined by magnitude of scaled differential expression. These figures have been filtered such that top 100 edges representing most differentially expressed node pairs are shown. (n) Bar chart showing quantification of Fig. 5o. Data are means ± s.e.m, P values calculated by two-sided *t*-test. n.s., no significance, \*\*\**P*<0.001, \*\*\*\**P*<0.0001, each sample was counted from three different slices.

**Supplementary Figure 6.**
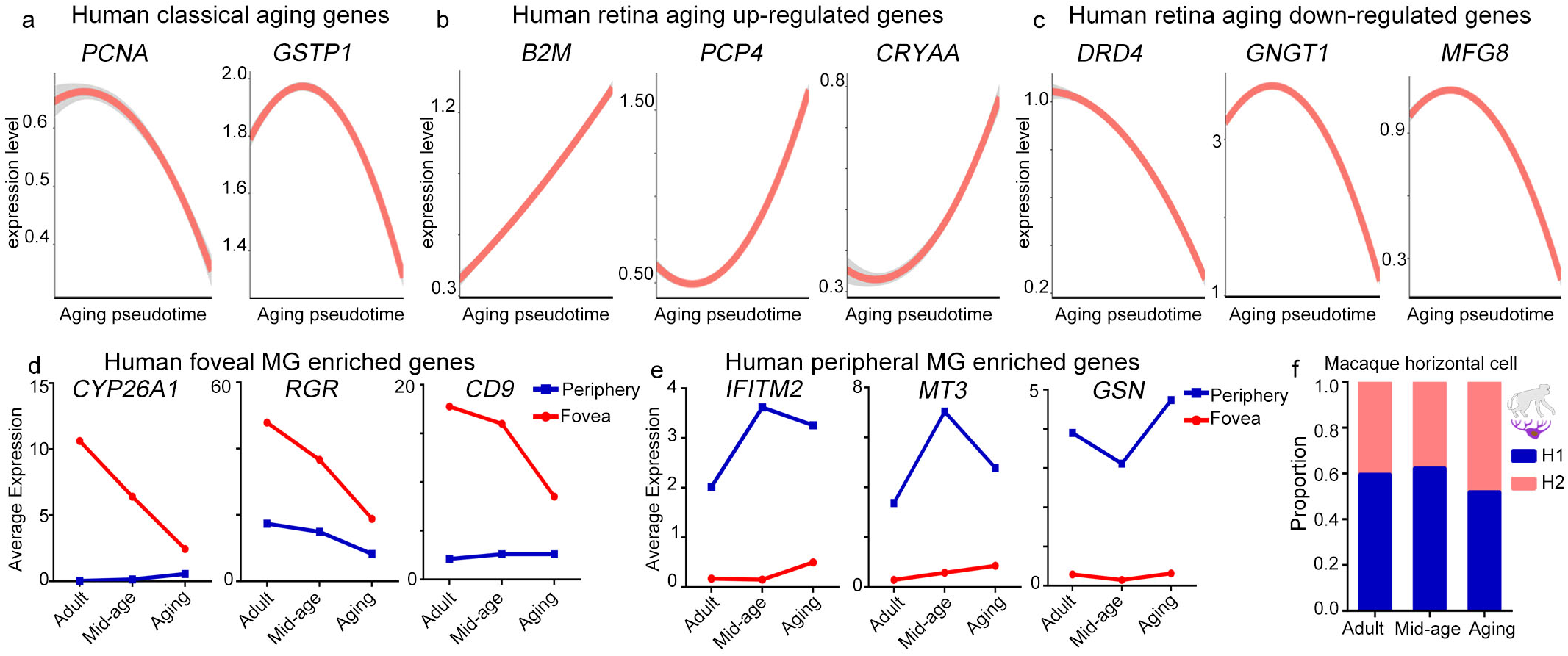
Trajectories of human retinal aging. (a) Trajectories of expression of classical human aging genes across human adult retina aging pseudotime. The shadow represents the confidence interval (95%) around the fitted curve. (b-c) Trajectories of expression of up-regulated (b) and down-regulated (c) genes across human adult retina aging pseudotime. The shadow represents the confidence interval (95%) around the fitted curve. (d-e) Average expression of genes enriched in foveal (d) and peripheral MGs (e). (f) Cell proportions of macaque H1 and H2 subclasses at different aging stages.

**Supplementary Figure 7.**
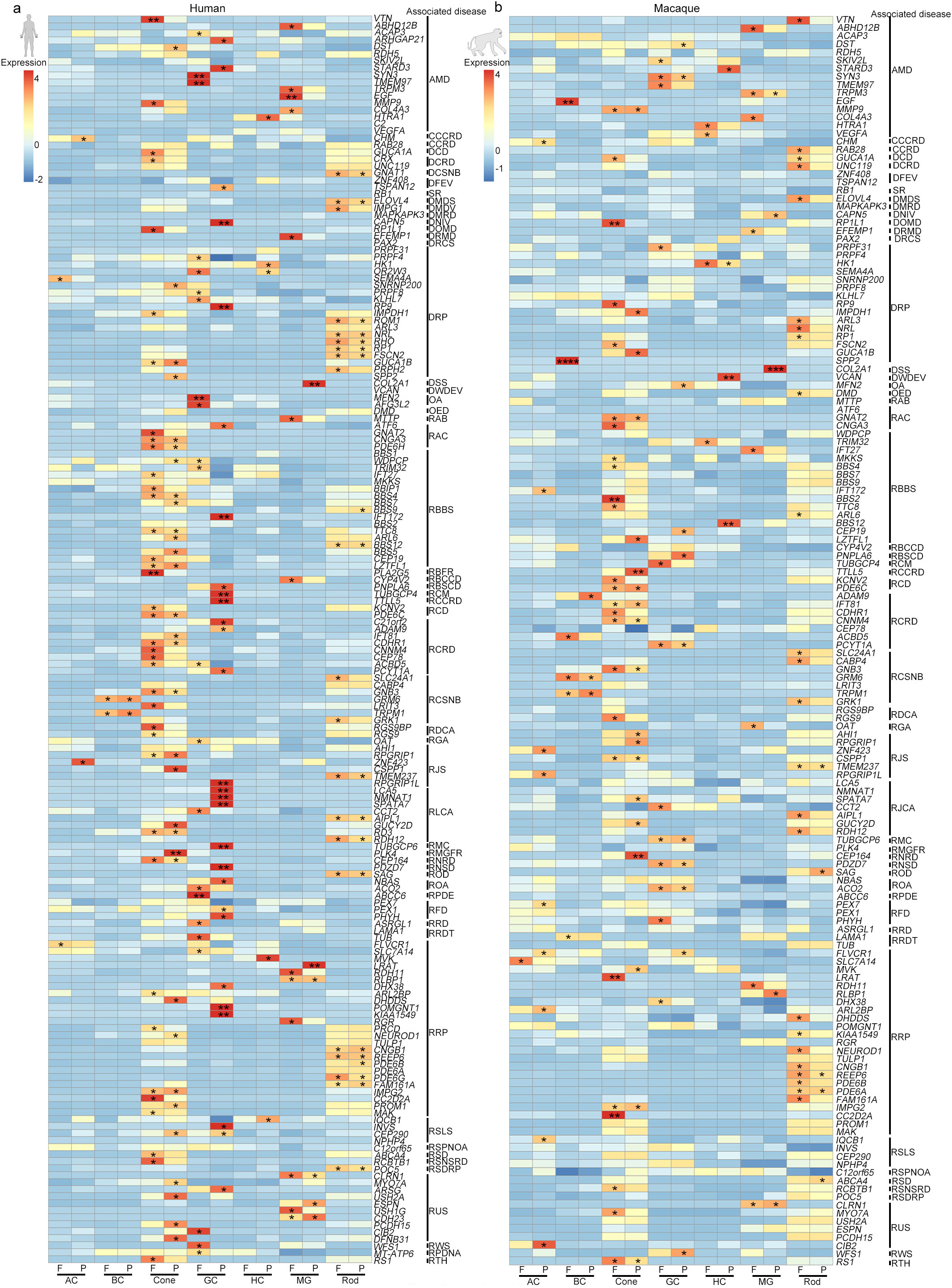
Expression patterns of genes in humans and macaques related to human retinal diseases. (a-b) Expression patterns of human eye disease-associated genes in human (a) and macaque (b) retinal cell types. AMD: age-related macular degeneration. CCCRD: choroideremia, cone and cone-rod dystrophy. CCRD: cone and cone-rod dystrophy. DCD: dominant cone dystrophy. DCRD: dominant cone-rod dystrophy. DCSNB: dominant congenital stationary night blindness. DFEV: dominant familial exudative vitreoretinopathy. SR: somatic retinoblastoma. DMDS: dominant macular dystrophy, Stargardt-like. DMDV: dominant macular dystrophy, vitelliform. DMRD: dominant Martinique retinal dystrophy and retinitis pigmentosa. DNIV: dominant neovascular inflammatory vitreoretinopathy. DOMD: dominant occult macular dystrophy. DRMD: dominant radial macular drusen. DRCS: dominant renal-coloboma syndrome. DRP: dominant retinitis pigmentosa. DSS: dominant Stickler syndrome. DWDEM: dominant Wagner disease and erosive vitreoretinopathy. OA: optic atrophy. OED: Oregon eye disease. RAB: recessive abetalipoproteinemia. RAC: recessive achromatopsia. RBBS: recessive Bardet Biedl syndrome. RBFR: recessive benign fleck retina. RBCCD: recessive Bietti crystalline corneoretinal dystrophy. RBSCD: recessive Boucher-Neuhauser syndrome with chorioretinal dystrophy. RCM: recessive chorioretinopathy and microcephaly. RCCRD: recessive cone and cone-rod dystrophy. RCD: recessive cone dystrophy. RCRD: recessive cone-rod dystrophy. RCSNB: recessive congenital stationary night blindness. RDCA: recessive delayed cone adaptation. RGA: recessive gyrate atrophy. RJS: recessive Jobert syndrome. RLCA: recessive Leber congenital amaurosis. RMC: recessive microcephaly with chorioretinopathy. RMGFR: recessive microcephaly, growth failure and retinopathy. RNRD: recessive nephronophthisis with retinal degeneration. RNSD: recessive non-syndromic deafness. ROD: recessive Oguchi disease. ROA: recessive optic atrophy. RPDE: recessive pseudoxanthoma elasticum. RFD: recessive refsum disease. RRD: recessive retinal disease. RRDT: recessive retinal dystrophy. RRP: recessive retinitis pigmentosa. RSLS: recessive Senior-Loken syndrome. RSPNOA: recessive spastic paraplegia, neuropathy, and optic atrophy. RSD: recessive Stargardt disease. RSNSRD: recessive syndromic and non-syndromic retinal dystrophy. RSDRP: recessive syndromic disease with retinitis pigmentosa. RUS: recessive Usher syndrome. RWS: recessive Wolfram syndrome. RPDNA: retinitis pigmentosa with developmental and neurological abnormalities. RTH: retinoschisis. *P* values were calculated by bootstrap hypothesis test, \**P* <0.05, ** *P* <0.01, *** *P* <0.001, \*\*\*\**P*<0.0001.

**Supplementary Table1. Retina sample information.** Spread sheet includes human and macaque adult retinal sampling information.

**Supplementary Table2. Top 100 DEGs of different cell types of human retina.** Spread sheet includes 100 marker genes of all 10 major cell types of human retina.

**Supplementary Table3. Top 100 DEGs of different cell types of macaque retina.** Spread sheet includes 100 marker genes of all nine major cell types of macaque retina.

**Supplementary Table 4. Up- and down-regulated genes of human retinal aging.** Spread sheets include 87 and 121 up- and down-regulated genes, respectively, when human retina age.

**Supplementary File 1**. Codes for bioinformatic analysis.

